# A zebrafish model for COVID-19 recapitulates olfactory and cardiovascular pathophysiologies caused by SARS-CoV-2

**DOI:** 10.1101/2020.11.06.368191

**Authors:** Aurora Kraus, Elisa Casadei, Mar Huertas, Chunyan Ye, Steven Bradfute, Pierre Boudinot, Jean-Pierre Levraud, Irene Salinas

## Abstract

The COVID-19 pandemic has prompted the search for animal models that recapitulate the pathophysiology observed in humans infected with SARS-CoV-2 and allow rapid and high throughput testing of drugs and vaccines. Exposure of larvae to SARS-CoV-2 Spike (S) receptor binding domain (RBD) recombinant protein was sufficient to elevate larval heart rate and treatment with captopril, an ACE inhibitor, reverted this effect. Intranasal administration of SARS-CoV-2 S RBD in adult zebrafish recombinant protein caused severe olfactory and mild renal histopathology. Zebrafish intranasally treated with SARS-CoV-2 S RBD became hyposmic within minutes and completely anosmic by 1 day to a broad-spectrum of odorants including bile acids and food. Single cell RNA-Seq of the adult zebrafish olfactory organ indicated widespread loss of expression of olfactory receptors as well as inflammatory responses in sustentacular, endothelial, and myeloid cell clusters. Exposure of wildtype zebrafish larvae to SARS-CoV-2 in water did not support active viral replication but caused a sustained inhibition of *ace2* expression, triggered type 1 cytokine responses and inhibited type 2 cytokine responses. Combined, our results establish adult and larval zebrafish as useful models to investigate pathophysiological effects of SARS-CoV-2 and perform pre-clinical drug testing and validation in an inexpensive, high throughput vertebrate model.

## Introduction

The current COVID-19 pandemic has infected 44.8 million people worldwide causing over 1 million global deaths as of October 21st 2020. Consequently, development of laboratory animal models that recapitulate the pathophysiology of SARS-CoV-2 infection in humans is of utmost importance to accelerate drug and vaccine testing. Thus far, the few lab compatible animal species that have been reported as naturally susceptible to SARS-CoV-2 include rhesus and cynomolgus macaques (Munster et al., 2020; Rockx et al., 2020), ferret (Kim et al., 2020), cat (Shi et al., 2020), and Syrian hamster (Chan et al., 2020). Mice, by contrast, are not spontaneously permissive to the virus, but mice expressing the human ACE2 receptor provide a useful animal model (Bao et al., 2020; Jia et al., 2020; Lutz et al., 2020). All these mammalian models have unique advantages and disadvantages for the study of immune responses to SARS-CoV-2 and other host-pathogen interactions but do not allow rapid, whole organismal, high throughput, and low-cost preclinical testing of drugs and immunotherapies.

As model vertebrates, zebrafish are permissive to human viral pathogens including Influenza A (Gabor et al., 2014), Herpes Simplex virus type 1 (Burgos et al., 2008), Chikungunya virus (Palha et al., 2013) and human noroviruses GI and GII (Van Dycke et al., 2019). Zebrafish offer many advantages over other animal models due to their high reproductive ability, rapid development, low maintenance costs, and small transparent bodies. Importantly, zebrafish olfactory, immune, and cardiovascular physiology share a significant degree of conservation with humans (Postlethwait et al., 2020; Saraiva et al.). Genetically, more than 80% of disease related genes have a zebrafish orthologue (Howe et al., 2013). The zebrafish innate immune system is already developed in the transparent larval stages and members of all major groups of mammalian cytokines have been identified in the zebrafish genome (Gomes and Mostowy, 2020; Zou and Secombes, 2016). Academic laboratories and the pharmaceutical industry use zebrafish larvae in preclinical studies for assessing efficacy and toxicity of candidate drugs for several diseases (Taylor et al., 2010). Zebrafish is a proposed model for COVID-19 and has recently been used in one vaccination study (Galindo-Villegas, 2020; Ventura-Fernandes et al., 2020). This study aims to elucidate the physiopathology of wildtype zebrafish in response to SARS-CoV-2.

SARS-CoV-2 infection causes a wide litany of symptoms, ranging from asymptomatic to mild or severe disease (Menni et al., 2020). Apart from respiratory symptoms, multi-organ pathologies are often reported with heterogeneous symptoms such as olfactory and taste loss, cardiac dysfunction, renal pathologies, neurological damage, muscle and joint pain, gastrointestinal symptoms, clotting disorders and others (Tabata et al., 2020; Wang et al., 2020a; Zhu et al., 2020). A longitudinal study associated disease severity and patient outcome with specific patterns of innate immune response misfiring in COVID-19 patients leading to catastrophic organ damage (Lucas et al., 2020). Specifically, severe disease patients display maladaptive cytokine responses characterized by high levels of pro-inflammatory cytokines such as IL6, TNFα and IL1β (Blanco-Melo et al., 2020; Hadjadj et al., 2020; Lucas et al., 2020; Del Valle et al., 2020), suppression of type I IFN responses (Blanco-Melo et al., 2020; Hadjadj et al., 2020; Mudd et al., 2020), and elevated type 2 cytokine levels (Lucas et al., 2020). Importantly, this cytokine pattern is in sharp contrast to that found in patients experiencing mild or moderate symptoms, who are able to control exacerbated type 1 and type 3 cytokine responses (Lucas et al., 2020).

SARS-CoV-2 enters the human host cells when SARS-CoV-2 Spike (S) protein receptor binding domain (RBD) binds to angiotensin-converting enzyme 2 (ACE2) on a permissive host cell, then a serine protease, such as TMPRSS2, cleaves the spike protein S1/S2 site to facilitate fusion of the virion with the host cell membrane (Hoffmann et al., 2020a, 2020b). ACE2 is expressed in many different cell types across many organs in the human body including lung, olfactory sustentacular cells, enterocytes, and endothelial cells (Albini et al., 2020; Lamers et al., 2020; Ziegler et al., 2020; Zou et al., 2020), explaining the widespread organ damage observed in patients infected with SARS-CoV-2 (Wadman et al., 2020). Several studies performed comparative analyses of the ACE2 amino acid sequences across vertebrate species and identified a high degree of conservation use used ACE2 homology to predict host susceptibility to SARS-CoV-2 infection (Damas et al., 2020; Wan et al., 2020). Ectotherms such as fish are predicted to have a low propensity for SARS-CoV-2 infection. However, such *in silico* predictions must be ascertained by performing infection experiments.

Binding of the spike protein from SARS-CoV, NL63, and SARS-CoV-2 results in rapid and local decreases in ACE2 protein (Garvin et al., 2020; Glowacka et al., 2010; Imai et al., 2005; Kuba et al., 2005). In turn, decreased ACE2 levels trigger negative regulation of renin-angiotensin system (RAS) and bradykinin pathways ultimately responsible for hypotension, vascular permeability, and multiorgan damage (Garvin et al., 2020; Patel et al., 2014). Thus, COVID-19 patients can experience a variety of cardiovascular presentations including myocardial injury, atrial and ventricular arrhythmias and cardiogenic shock (Babapoor-farrokhran et al., 2020; Guzik et al., 2020; Kochi et al., 2020; Kuck, 2020; Libby, 2020). To restore balance to the RAS pathway and ameliorate cardiac symptoms associated with decreased ACE2, the use of ACE inhibitors is being considered as a therapeutic intervention in COVID-19 patients (Lopes et al., 2020). Importantly, drugs currently used to treat the COVID-19 can be pro-arrhythmic and therefore there is a need to incorporate cardiovascular damage into the list of targets of therapeutic interventions in COVID-19 and for models that replicate human cardia physiology (Kochi et al., 2020).

A hallmark of SARS-CoV-2 infection is acute loss of smell (Cooper et al., 2020). Viral-induced anosmia is not unique to SARS-CoV-2 infections since viruses such as rhinoviruses, influenza, parainfluenza and coronaviruses are known to be the main cause of olfactory deficits in humans (Suzuki et al., 2007; Imam et al., 2020). In mice and humans, *ace2* expression is detected in sustentacular cells, olfactory stem cells known as horizontal and globose basal cells in the olfactory epithelium, and vascular cells (pericytes) in the olfactory bulb (Brann et al., 2020). Olfactory dysfunction in COVID-19 patients may be caused by several mechanisms such as inflammation instigated by infection of cells in the olfactory epithelium altering olfactory sensory neuron’s (OSN) function and environment, as well as direct viral infection of sustentacular cells, olfactory stem cells, and vascular cells wreaking havoc (Cooper et al., 2020). Of note, sterile induction of type I IFN in the mouse olfactory epithelium leads to widespread loss of olfactory receptor expression and COVID-19 patients have increased pro-inflammatory cytokine levels in their olfactory epithelium (Rodriguez et al., 2020; Torabi et al., 2020). Still, experimental data is needed to elucidate the underlying mechanisms of loss of smell in COVID-19 patients.

The present study reports for the first time that zebrafish larvae exposed to SARS-CoV-2 appear to mount innate immune responses that resemble cytokine responses of mild COVID-19 patients. Recombinant SARS-CoV-2 S RBD is sufficient to cause olfactory, renal and cardiovascular pathologies in larvae and adult zebrafish. We also identify potential mechanisms of SARS-CoV-2 induced anosmia by scRNA-Seq. Our findings support the use of zebrafish as a novel vertebrate model to elucidate SARS-CoV-2 pathophysiology and to screen drugs and other therapies targeting COVID-19.

## Results

### Phylogenetic analyses of ACE2 molecules in vertebrates

Comparative analysis of ACE2 molecules in vertebrates indicated that ACE2 molecules are well conserved in vertebrates with a 72%-73% similarity and 57.5%-58% identity between zebrafish ACE2 and human ACE2, respectively (Table S1). Examination of ACE2 amino acid motifs in the region involved in binding SARS-CoV-2 S protein revealed zebrafish ACE2 has 50%/64% sequence similarity with the corresponding human ACE2 region compared to 71%/78% in macaques ACE2 or 57%/71% in ferret ACE2 (Table S2) (Liu et al., 2020). Combined, these data agree with previous reports and reveal that key amino acids involved in the interactions between human ACE2 and SARS-CoV-2 S protein are not conserved in zebrafish (Fu et al., 2020; Postlethwait et al., 2020).

### SARS-CoV-2 Spike (S) Receptor Binding Domain (SARS-CoV-2 S RBD) elevates heart rate in zebrafish larvae

Systemic injection of recombinant SARS-CoV-2 protein into adult zebrafish has been shown to induce some toxicity (Ventura-Fernandes et al., 2020). In order to determine whether recombinant SARS-CoV-2 S RBD protein causes inflammatory responses in zebrafish larvae, we exposed 5 dpf larvae to SARS-CoV-2 S RBD recombinant protein for 3 hours (h) and measured cytokine responses by qPCR. As shown in Figure 1A, 3 h immersion with SARS-CoV-2 S RBD protein induced a significant downregulation in *ifnphi1* expression and significant increase in expression of *ccl20a*.*3*, a pro-inflammatory chemokine. No changes in *ace2, tnfα, il1b* and *il17a/f3* expression were observed (Figure 1A). These results indicate that rapid immune responses occur in zebrafish larvae exposed to SARS-CoV-2 S RBD. We next evaluated the effects of SARS-CoV-2 RBD S on zebrafish larva heart function to validate zebrafish larvae as a model for COVID-19 cardiac manifestations. We immersed 7- and 5-days post fertilization (dpf) zebrafish larvae with SARS-CoV-2 S RBD, or with vehicle, and measured heart rate after 3 h. As shown in Figure 1B, 7 dpf and 5 dpf zebrafish treated with SARS-CoV-2 S RBD had significantly higher heart rates compared to vehicle treated controls. As a positive control for the recombinant protein, we used animals treated with the same dose of recombinant infectious hematopoietic necrosis virus (IHNV) glycoprotein (r-IHNVg), a rhabdovirus known to cause severe endothelial damage in zebrafish (Ludwig et al., 2011). r-IHNVg caused a severe decrease in larval zebrafish heart rate compared to control treated animals (Figure 1B-C). To determine if increased heart rate induced by SARS-CoV-2 S RBD was dependent on Ace2 binding, we co-incubated 5 dpf larvae with captopril and reverted SARS-CoV-2 S RBD induced heart dysfunction. Captopril had no effect on r-IHNVg induced bradycardia (Figure 1B). Ventricular trace analyses showed marked differences in rhythm patterns in each treatment group (Figure 1D). Importantly, the captopril and SARS-CoV-2 S RBD treated animals, despite having similar heart rates to those of the vehicle treated controls, displayed a unique ventricular trace pattern, warranting future studies regarding the potential cardioprotective role of captopril in COVID-19. Combined, these results indicate that zebrafish exposed to SARS-CoV-2 S RBD protein experience tachycardia and suggest that zebrafish larvae constitute a valuable pre-clinical model to test the effects of drugs for COVID-19 on cardiac activity *in vivo*.

**Figure 1:**
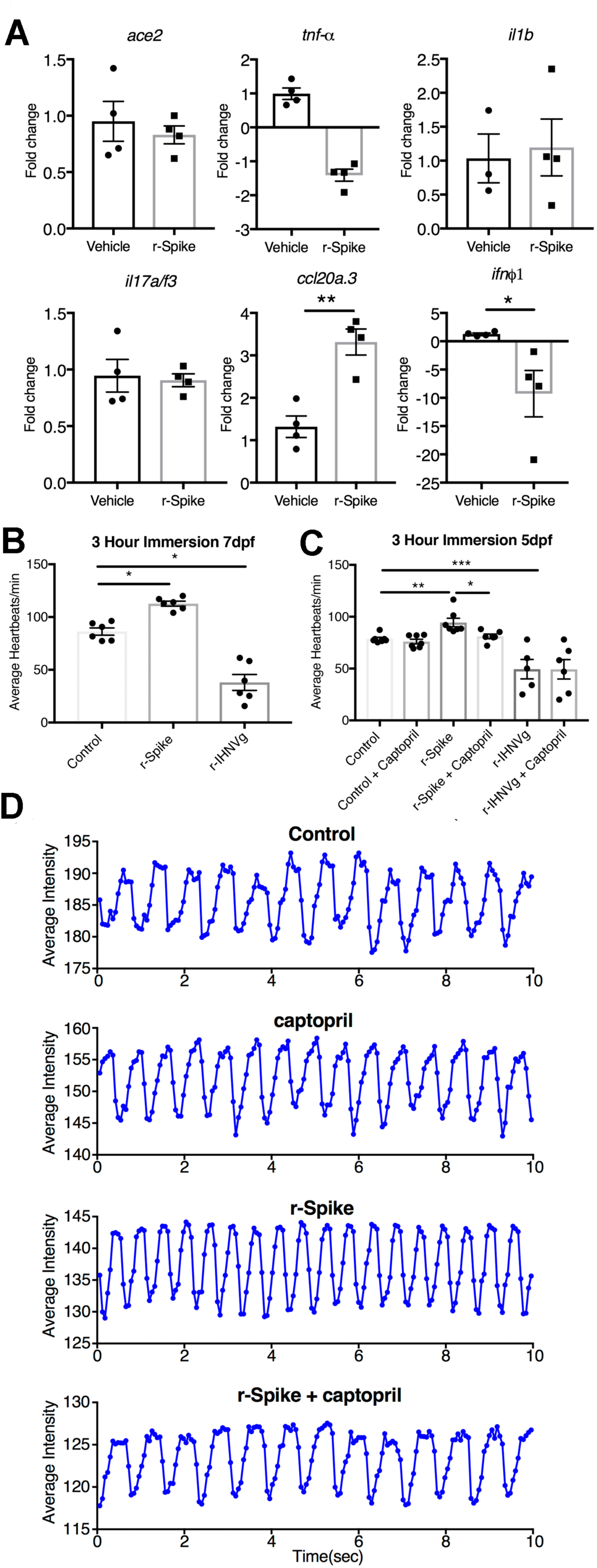
SARS-CoV-2 S RBD induces immune responses and elevates heart rate of larval zebrafish. (A) Changes in gene expression of *ace2, tnf-α, il1β, il17a/f3, ccl20a*.*3*, and *ifnphi1* in 5 dpf zebrafish larvae in responses to SARS-CoV-2 S RBD. Animals were exposed to SARS-CoV-2 S RBD protein (r-Spike, 2 ng/ml) for 3 h at 28.5°C or vehicle. Changes in gene expression were measured by RT-qPCR using *rps11* as the house-keeping gene. Each data point represents a pool of 4 larvae/well. Data are expressed fold-change compared to vehicle controls using the Pffafl method. (B) Average heart beat per minute of 7 dpf (n=6) zebrafish larvae after 3 h of incubation with vehicle, 2 ng/ml r-Spike, or 2 ng/ml r-IHNVg. (C) Average zebrafish heart beats per minute in 5 dpf zebrafish larvae (n=8) after 3 h of incubation with vehicle, 2 ng/ml r-Spike, or 2 ng/ml r-IHNVg with and without treatment with 12mM of captopril. Heart beats were recorded for 3 min at 20□ under a Nikon Ti microscope and videos were processed with ImageJ Time Series Analyzer V3. Data were analyzed by one-way ANOVA with Tukey’s multiple comparisons post hoc. Results are representative of two independent experiments. (D) Representative ventricular traces show the average intensity in region of interest (heart ventricle) in 5 dpf zebrafish treated for 3 h with vehicle, or 2 ng/ml r-Spike, 12 mM captopril or 12mM captopril and 2 ng/ml r-Spike. Traces are representative of one animal per treatment (n=8 per treatment group). * *P-*value<0.05; ** *P-*value<0.01 *** *P-*value<0.001.

### Intranasal delivery of SARS-CoV-2 S RBD causes olfactory and renal damage in adult zebrafish

Anosmia is one of the earliest manifestations of SARS-CoV-2 infection in humans (Cooper et al., 2020). We have previously shown that IHNV glycoprotein protein is sufficient to induce rapid nasal immune responses as well as neuronal activation in teleost fish (Sepahi et al., 2019). Thus, we hypothesized that recombinant SARS-CoV-2 S RBD protein may be sufficient to damage the olfactory epithelium of zebrafish. In order to test this hypothesis, we delivered recombinant SARS-CoV-2 S RDB to the nasal cavity of adult AB zebrafish and sampled the olfactory organ 3 h, 1 day (d), 3 d and 5 d later. Negative controls consisted of animals that received vehicle only (PBS) into the nasal cavity. As shown in Figure 2, SARS-CoV-2 S RBD was sufficient to cause severe damage to the zebrafish olfactory organ at all time points examined. Tissue damage was characterized by presence of edema and endothelial inflammation in the lamina propria as early as 3 h post-treatment (Figure 2A-C and G). Presence of hemorrhages at the tip and base of the olfactory lamellae was observed 1 d post-treatment (Figure 2D). Loss of the epithelial mosaic structure characteristic of the teleost olfactory epithelium was observed on days 1, 3 and 5 post-treatment (Figure 2E). By day 5, loss of entire apical lamellar areas due to severe necrosis was observed in the olfactory lamellae of all treated animals compared to controls (Figure 2F). Significant loss of olfactory cilia was recorded in all animals treated with SARS-CoV-2 S RBD at all time points (Figure 2H). These results indicate the SARS-CoV-2 S RBD is sufficient to cause inflammation, edema, hemorrhages, ciliary loss, and necrosis in the olfactory organ of zebrafish. Hence, olfactory damage can be caused by indirect mechanisms and in the absence of active SARS-CoV-2 replication in this tissue.

**Figure 2:**
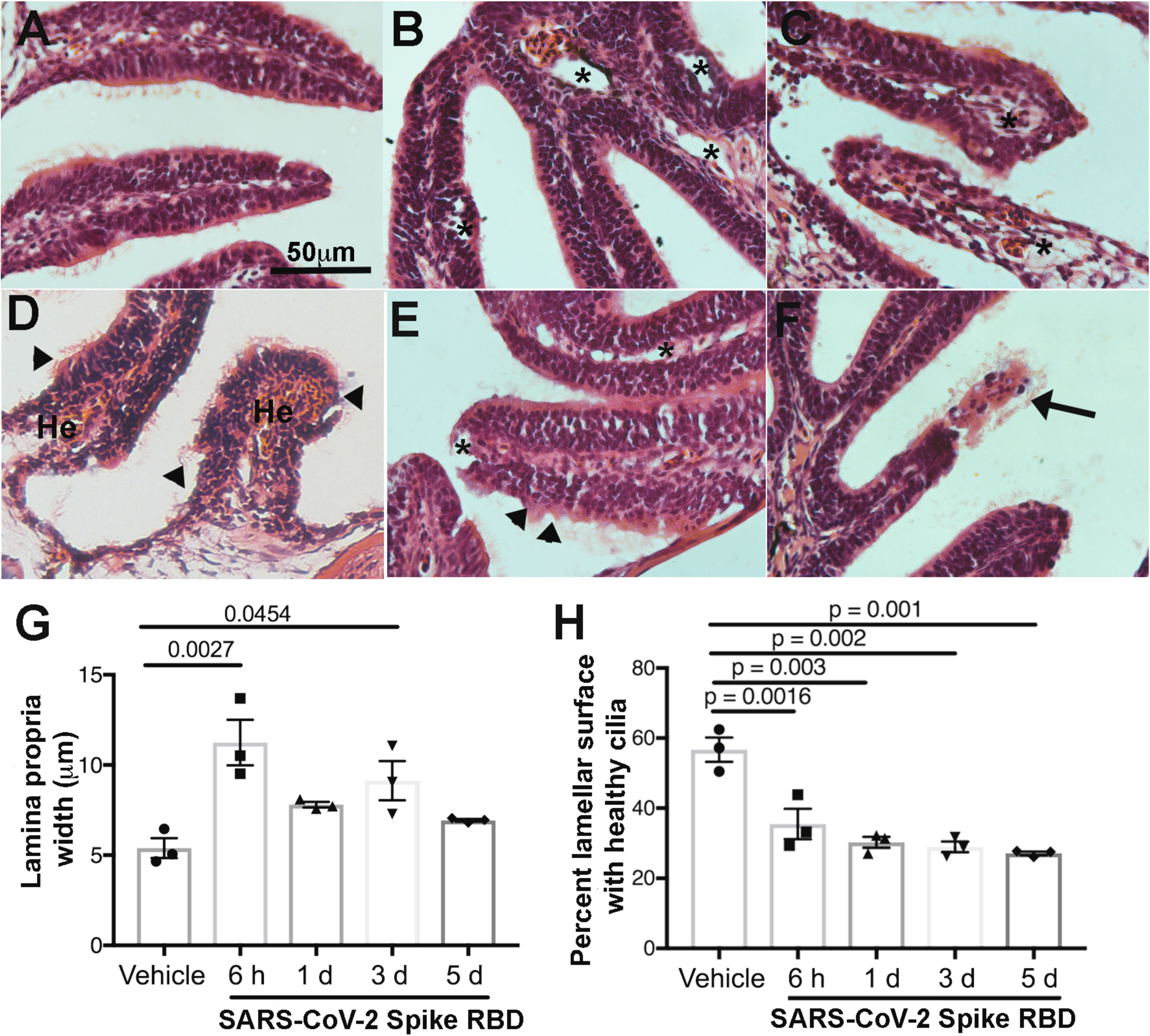
Intranasal delivery of SARS-CoV-2 S RDB protein causes severe olfactory damage in adult zebrafish. H&E stains of zebrafish olfactory organ paraffin sections. (A) Control vehicle treated olfactory organ. (B-C) 3 h; (D) 1 d; (E) 3 d; (F) 5 days post intranasal delivery of SARS-CoV-2 S RDB protein (r-Spike). Images are representative of n=10 sections per animal and n=3 fish per experimental group. Asterisk denotes edemic spaces in the olfactory lamellae, mostly in the lamina propria. He: hemorrhages. Arrow heads: apical loss of olfactory sensory neurons. Black Arrow: Necrotic lamella. Image analyses in G and H were performed in ImageJ. Differences were considered statistically significant when *P-*value<0.05.

Toxicity effects of intranasal SARS-CoV-2 S RBD delivery were also evaluated in distant tissues such as the kidney, a target organ of SARS-CoV-2. Acute kidney injury (AKI) incidence varies from 0.9% to 29% in COVID-19 patients (Su et al., 2020). Renal damage, especially AKI, is also common in patients with RAS dysfunction such as diabetic patients who suffer from hypertension (Ribeiro-Oliveira et al., 2008; Advani, 2020). A recent study in zebrafish injected with the N-terminal part of SARS-CoV-2 S protein reported inflammation and damage in several tissues of adult zebrafish including kidney 7 and 14 d post-injection (Ventura-Fernandes et al., 2020). In the present study, histological examination of the head-kidney of zebrafish who received SARS-CoV-2 S RBD intranasally revealed renal tubule pathology characteristic of AKI 3 h post-treatment (Figure S1). Pathology was not as severe at later time points, but vacuolation of the renal tubule epithelium was still visible 5 days post-treatment (Figure S1). We did not observe signs of glomerulopathology in treated animals compared to controls. Together, these results indicate that intranasal delivery of SARS-CoV-2 S RBD is sufficient to cause nephropathy in adult zebrafish but that pathology is not as severe as when the protein is delivered by injection.

### Intranasal delivery of SARS-CoV-2 S RBD causes anosmia in adult zebrafish

Adult zebrafish exposed to SARS-CoV-2 S RBD had a significant reduction of olfactory responses to food extracts of ∼50% of preexposure olfaction within minutes as measured by electro-olfactogram (EOG), indicating an instant effect of the protein on olfactory function (Fig. 3A). Reduction of olfaction was sustained for least one hour of recording, but the olfactory organ remained still semi functional. Zebrafish treated with PBS never lost olfaction at any time point. To further quantify the degree of olfactory reduction due to SARS-CoV-2 S RBD, we took advantage of the two easily accessible and isolated olfactory chambers present in zebrafish. We exposed one naris to the SARS-CoV-2 S RBD protein and the other naris, from the same animal, to PBS and waited 3 h or 1d before measuring olfaction by EOG. At 3 h we observed a 37-70% reduction in food and bile olfactory responses between the treated and untreated naris and a complete loss of olfactory function to both odorants 1 d post-treatment (Fig. 3B). The reduction of olfactory sensitivity for food extract was smaller than that found for bile, probably due to the lower number of OSNs involved in bile acid detection compared to amino acids found in food (Hansen et al., 2003). Our results indicate that SARS-CoV-2 S RBD-induced-anosmia is not specific for a subset of OSNs, since both food extracts and bile olfactory signals were suppressed in SARS-CoV-2 S RBD treated zebrafish. This fact, together with the 1d time to develop complete anosmia and disrupt olfactory epithelial structure, support the hypothesis that SARS-CoV-2 S RBD damage may occur first on sustentacular cells, with subsequent impacts on OSN viability and function.

**Figure 3:**
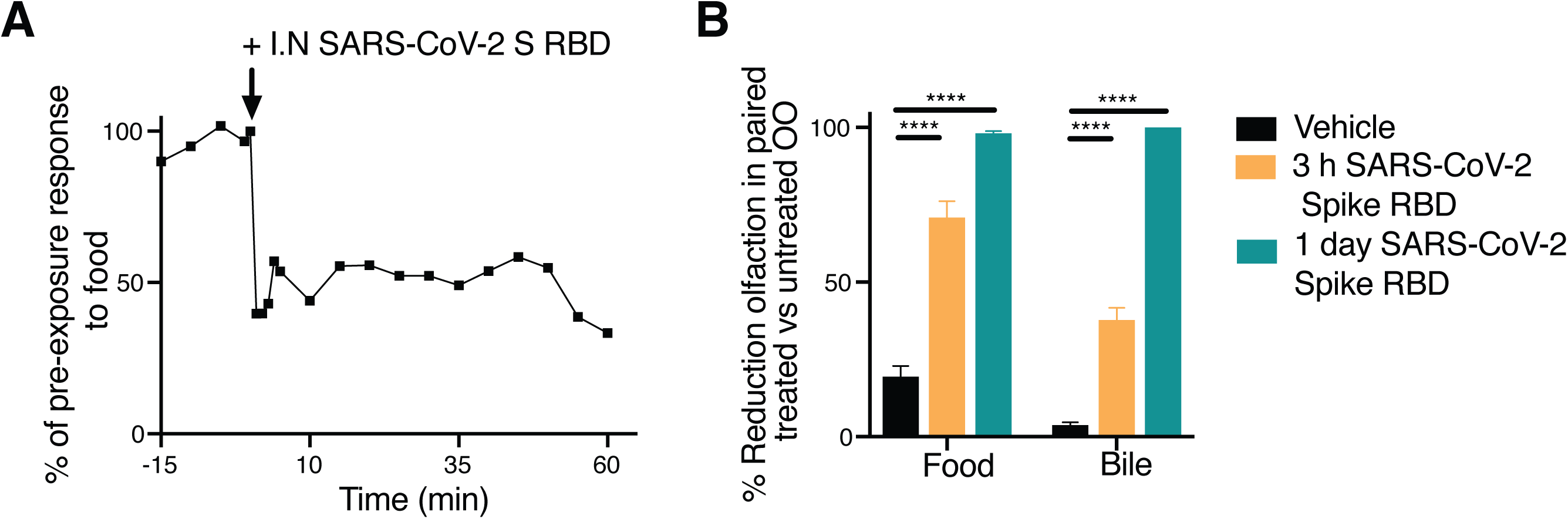
Intranasal delivery of SARS-CoV-2 S RDB protein causes anosmia in adult zebrafish. (A) Representative EOG of adult zebrafish showing olfactory responses to food over time before and after addition of SARS-CoV-2 S RDB protein to the naris. (B) Mean percentage loss of olfactory detection of food or bile salts measured by EOG recording. Adult zebrafish (n=4) treated with SARS-CoV-2 S RDB protein in one naris or vehicle control in the other naris for 3 h or 1 d after which EOG recorded as animal exposed to food odorant or bile in both nares. Percent reduction calculated as percent of the SARS-CoV-2 S RDB treated naris’ EOG response to vehicle treated naris’ EOG response. Statistical analyses are ANOVA with Tukey’s post hoc multiple comparison **** *P*-value < 0.0001.

### Single-cell analysis of the zebrafish olfactory organ

To understand the impact of SARS-CoV-2 S RBD on zebrafish OO, we performed single cell RNA-Seq (scRNA-Seq) of adult zebrafish olfactory organ from control or SARS-CoV-2 S RBD intranasally-treated fish. scRNA-Seq was performed after 3 h or 3 d of intranasal administration with SARS-CoV-2 S RBD. These timepoints were selected based on the histological findings of Figure 2. Data from the treated time point was integrated with that from vehicle treated controls in Seurat and then clustered using 30 significant principal components (Figure 4A-B)(Butler et al., 2018). We identified a total of 8 neuronal cell clusters (NC), 5 sustentacular cell (SC) clusters, 3 endothelial cell (EC) clusters and 7 leucocyte (lymphoid and myeloid) clusters (Figure 4A-B).

**Figure 4:**
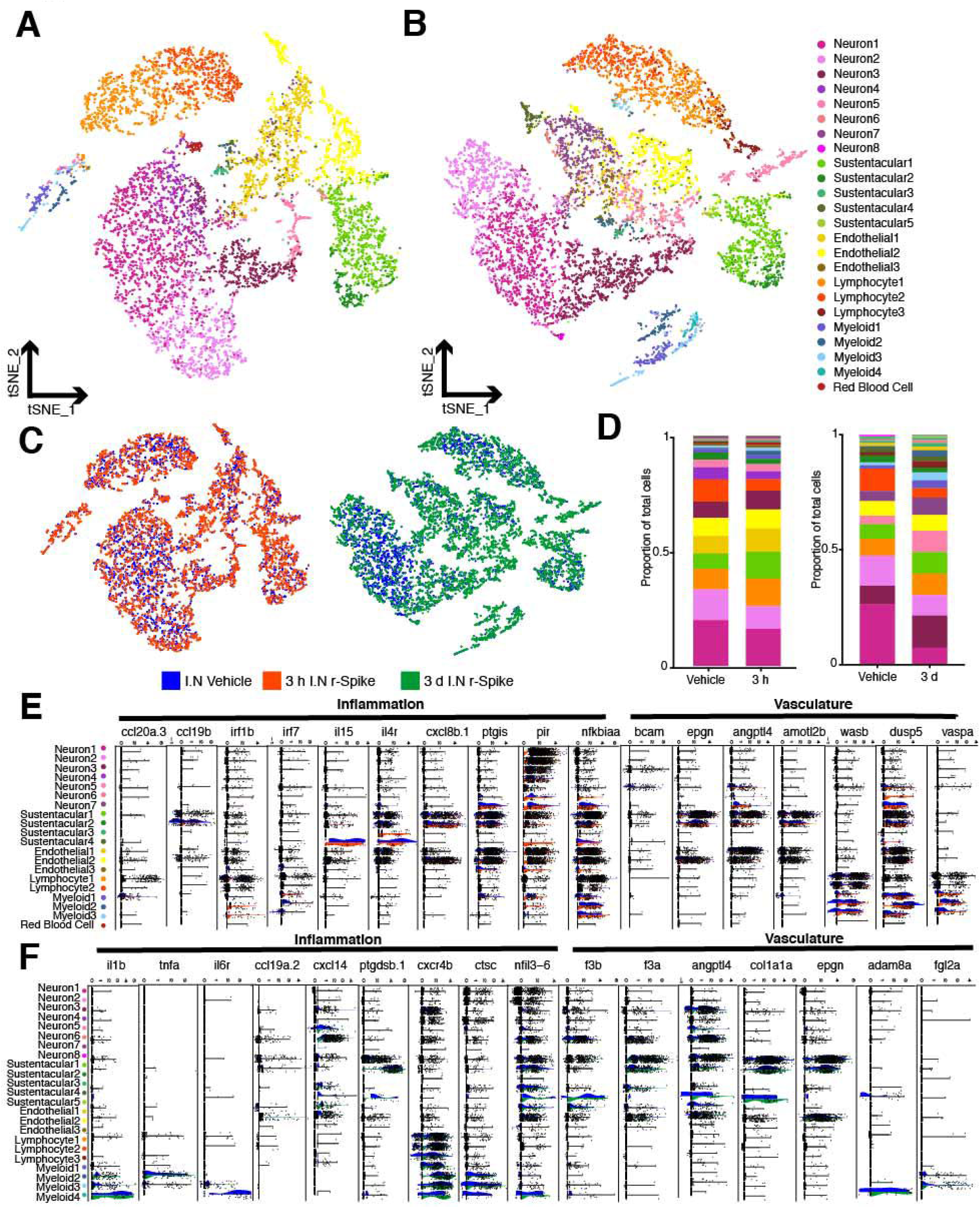
ScRNA-Seq of zebrafish olfactory organ reveals changes in cellular landscape due to intranasal SARS-CoV-2 S RBD protein delivery. (A-B) Cellular composition of the zebrafish olfactory organ visualized using t-distributed stochastic neighbor embedding (tSNE) of vehicle treated, 3 h and 3 d post-intranasal delivery of SARS-CoV-2 S RBD (r-Spike). Individual single-cell transcriptomes were colored according to cluster identity. tSNE plots were generated in Seurat and identified 20 and 22 distinct cell clusters, respectively. (C) tSNE plots of zebrafish olfactory organ cell suspensions colored according to treatment. (D) Bar plots showing the proportion of cells belonging to each cluster in each treatment group. (E-F) Violin plots showing expression of selected markers of inflammation and vasculature remodeling in the 3 h SARS-CoV-2 S RBD (orange) versus vehicle treated (blue) and 3 d SARS-CoV-2 S RBD (green) versus vehicle treated (blue) zebrafish olfactory organ obtained by scRNA-Seq. All represented genes were differentially regulated between treatments (adjusted *P*-value <0.05).

Of the 8 NCs, Neuron1 and Neuron2 corresponded to mature OSNs. Neuron1 expressed markers of ciliated OSNs (*ompa* and *ompb*) in addition to several olfactory receptor (OR) genes (Buiakova et al., 1996). Neuron2, on the other hand, expressed markers of microvillus OSNs (*trpc2b, s100z*, and *gnao)* and many vomeronasal receptors (VR) such as *v2rh32* as well as OR gene (Figure S2) (Kraemer et al., 2008). Neuronal clusters 3-8 included OSNs in different differentiation stages, from progenitor-like neurons to differentiating intermediate neuronal precursors (Yu and Wu, 2017). For instance, expression of multiple high mobility group transcripts (*hmga1a*, and *hmgb2b2)* typically found in neuronal precursor cells is inverse to expression of the transcription factor family of sox genes (*sox4* and *sox11*) associated with committing to differentiation to mature neurons across the neuronal subsets (Bergsland et al., 2006; Giancotti et al., 2018). The Neuron3 cluster expressed both *ompa* and OR genes as well as markers of neuronal differentiation and plasticity such as *neurod1, neurod4, gap43, sox11*, and *sox4* suggesting commitment to becoming mature ciliated OSNs. The cluster Neuron4, only found at 3 h, was transcriptionally similar to the Neuron3 cluster expressing *hmga1a*, and *hmgb2b2*, but lacked markers of differentiation (*gap43* and *sox11)* indicating a closer neuronal progenitor state. The clusters Neuron5 and Neuron6 were the closest to the progenitor state with expression of basic helix-loop-helix transcription factors (i.e. *bhlhe23*), associated with exit from the cell cycle and progenitor state, as well as cell cycle genes (Borders et al., 2007). Clusters Neuron7 and Neuron8 expressed cell cycle and early neuronal progenitor markers as well as *tmprss13b* and *tmprss4a*, however the majority of cluster identifying genes in these two clusters are undescribed (Figure S2).

SCs are supporting cells that exist in the neuroepithelium around OSNs and in humans, they express ACE2 (Bryche et al., 2020). SCs in the olfactory epithelium can directly arise from horizontal basal cells (HBCs) (Yu and Wu, 2017). We found 5 clusters of SCs in our datasets. Sustentacular clusters 1 and 2 had similar expressing markers to those of HBCs (*krt5*) and other basal cells (*lamc2* and *col1a1b*) and therefore may represent olfactory basal cells (Chen et al., 2017; Peterson et al., 2019). Further, these clusters expressed immune genes such as *ptgs* and *ccl25*, and HBCs are known to express immune factors in models of olfactory epithelium damage (Chen et al., 2019). Interestingly, in addition to *sox2* expression, Sustentacular3 cluster expressed *atoh1*, a basic helix loop helix gene important in hair cell regeneration (Bermingham-McDonogh and Reh, 2011). Cluster Sustentacular4 expressed cytokines *il15* and *il17a/f3* as well as *ace*. The last SC cluster was only found in the 3 d time point and markers suggest it is a subpopulation closely related to the Sustentacular4 cluster (Figure S2).

We identified three clusters of ECs that all express *tmprss13* and *tmprss4*. While we did not detect *ace2* expression in any cell clusters, we detected *ace2* mRNA in adult zebrafish olfactory organ, and at low levels in the olfactory bulb (Figure S3). *ace2* expression levels have previously shown to be low in neuronal tissues and therefore may be hard to detect by scRNA-Seq (Song et al., 2020). Endothelial1 and 2 clusters expressed the endothelial markers *sox7* and *tmp4a* (Yao et al., 2019). All three clusters broadly expressed genes associated with tight junctions (*tjp3, jupa, ppl, cldne*, and *cgnl1*) as well as many keratin genes (Figure S2). Interestingly, Endothelial3 cluster also expressed the calcium channel *trpv6* and a slew of non-annotated genes.

There are copious amounts of immune cells in the teleost olfactory organ (Mellert et al., 1992; Sepahi et al., 2016), and although our method of generating single cell suspensions for scRNA-Seq was optimized for neuronal cell viability, we captured small groups of both myeloid and lymphoid cells in our samples. Lymphocyte1 cells expressed classical T cell markers (*cd8, cd4*, and *zap70*) as well as *ccr9*, a mucosal homing chemokine receptor, and novel immune type receptor genes (*nitrs*) thought to be natural killer cell receptors (Papadakis et al., 2003; Wei et al., 2007). Therefore, we propose that lymphocyte1 are a type of innate-like NKT cells. Lymphocyte2 cluster cells expressed the transcription factor *foxp3b* found in regulatory T cells (T_regs_). The 3 d time point revealed a third lymphocyte cluster with classical T cell identity. The myeloid cell clusters 1-3 expressed MHC II complex genes as well as *mpeg1*.*1*, a marker of zebrafish macrophages (Ellett et al., 2011). Myeloid4 cluster expressed *mpx*, a marker for neutrophils, and no antigen presentation genes (Yoo and Huttenlocher, 2011). Each macrophage-like myeloid cluster expressed unique cytokines contributing to their separate clustering --Myeloid1 expressed *il12* and *il22*, Myeloid2 expressed *il1fma* and *il10ra*, and Myeloid3 expressed *il6r* and *il1b*. Together, these data indicate that the zebrafish olfactory organ is a heterogeneous neuroepithelium with diverse immune cell subsets.

### Intranasal delivery of SARS-CoV-2 S RBD induces inflammatory responses and widespread loss of olfactory receptor expression in adult zebrafish olfactory organ

The cellular landscape of the zebrafish olfactory epithelium was affected by SARS-CoV-2 S RBD treatment and time (Figure 4A-D). This was especially evident in the proportions of neuronal cell types 3 d post-treatment when the proportion of mature, *omp*^+^ ciliated OSN was much lower compared to controls and the 3 h treated group. In contrast, neuronal progenitors expressing cell cycle markers (*aubk, ecrg4*, and *mki67*) and neuronal differentiation and plasticity markers (*neurod1, neurod4, gap43, sox11*, and *sox4*) were expanded 3 d post-treatment (Figure 4B-C). Further, we detected a noticeable decrease in the proportion of cells belonging to the Lymphocyte2 cluster, a cluster that expressed markers of Treg cells (*foxp3b*) 3 h post intranasal delivery of recombinant SARS-CoV-2 S RBD but this change was not noticeable at day 3 (Figure 4C). At 3 d, we observed a third lymphocyte cluster, not detected at 3 h, highly expressing the TCR subunit *zap70* as well as plac8 onzin related protein (*ponzr1*), a molecule that has immunoregulatory roles during Th1 type immune responses in mammals (Figure S2) (Slade et al., 2020). These results indicate important cellular responses in neuronal and immune cell subsets of the zebrafish olfactory organ in response to SARS-CoV-2 S RBD.

The transcriptional landscape of the zebrafish olfactory epithelium was affected by SARS-CoV-2 S RBD treatment (Figure 4E-F). By 3 h post intranasal delivery of recombinant SARS-CoV-2 S RBD, SCs and ECs showed significant increased expression of cytokine genes (*il15, il4r*, and *ccl19b*) and prostaglandin I2 synthase (*ptgis*), while myeloid clusters upregulated expression of the interferon gene *irf1b* (Figure 4E). Strikingly, NF-kB inhibitor alpha (*nfkbiaa*) was differentially expressed across almost all cell types including progenitor-like neuronal clusters at 3 h relative to vehicle control (Figure 4E). By 3 d post-treatment with recombinant SARS-CoV-2 S RBD, myeloid clusters expressed inflammatory markers (*il1b* and *tnfa*) and showed a significant differential upregulation of the chemokine receptor *cxcr4b* suggesting perhaps trafficking into the olfactory organ from other body sites (Figure 4F). At 3 h post intranasal delivery of SARS-CoV-2 S RBD, differential expression analysis showed changes in genes related to endothelial integrity including angiopoietin-like 4 (*angptl4*), angiomotin (*amotl2b*), and Wiskott-Aldrich syndrome protein (*wasb*) as well as other genes associated with vasoconstriction (*dusp5* and *vaspa*) (Figure 4E) (Li et al., 2015; Wagener et al., 2016; Zheng et al., 2009). By 3 d after intranasal SARS-CoV-2 S RBD, endothelial clusters showed significantly increased expression of clotting factors (*f3b* and *f3a*) relative to vehicle treated control and continued upregulation of *angptl4* expression was detected in several EC and SC clusters (Figure 4F). Thus, this data suggested that inflammatory responses and endothelial disruption are both hallmarks of SARS-CoV-2 induced damage in the olfactory epithelium.

Differential expression analysis revealed that while some changes in ORs expression occurred 3 h post-treatment in neuronal progenitor clusters, many ORs and TAARS genes were significantly differentially downregulated in mature neuronal clusters 1 and 2 compared to vehicle treated control (Figure 5A-B). In accord, markers of mature ciliated and microvillus OSNs (*ompb* and *v2rh32*, respectively) were significantly downregulated at both 3 h and 3 days after intranasal SARS-CoV-2 S RBD delivery (Figure 4A-D). Gene ontology (GO) analyses of significantly down or up regulated genes from all neuronal clusters at 3 h or 3 d after intranasal SARS-CoV-2 S RBD compared to vehicle treated controls was done using ShinyGO v0.61 and Metascape to gather an image of how OSNs as a collective are responding to the viral moiety (Figure 5E-H) (Ge et al., 2020; Zhou et al., 2019). At 3 h after intranasal SARS-CoV-2 S RBD delivery, upregulated genes in the olfactory neuronal clusters were enriched in regulation of cell differentiation and sensory perception of smell (Figure 5E). By 3 d, upregulated genes identified in olfactory neuronal clusters were enriched in processes such as neuron differentiation, sensory system development, and sensory organ morphogenesis, whereas downregulated genes belonged to sensory perception of smell, detection of chemical stimulus and GPCR signaling pathway (Figure 5F). Functional enrichment analyses in Metascape showed that the top non-redundant enriched clusters in both upregulated and downregulated genes in zebrafish OSNs 3 h post-treatment was sensory perception of smell (Figure 5G). The same was true 3 days post-treatment, but processes such as regeneration, neuron development, neuron fate commitment and the p53 signaling pathway were also enriched within the upregulated genes (Figure 5H). Combined, these results suggest that presence of SARS-CoV-2 S RBD in the olfactory organ instigates harmful effects on OSNs within hours and that the magnitude of the OSN damage increases by 3 days post-treatment. Further, these analyses indicate that neuronal regeneration and differentiation processes were initiated by day 3 in order to begin repair of olfactory damage.

**Figure 5.**
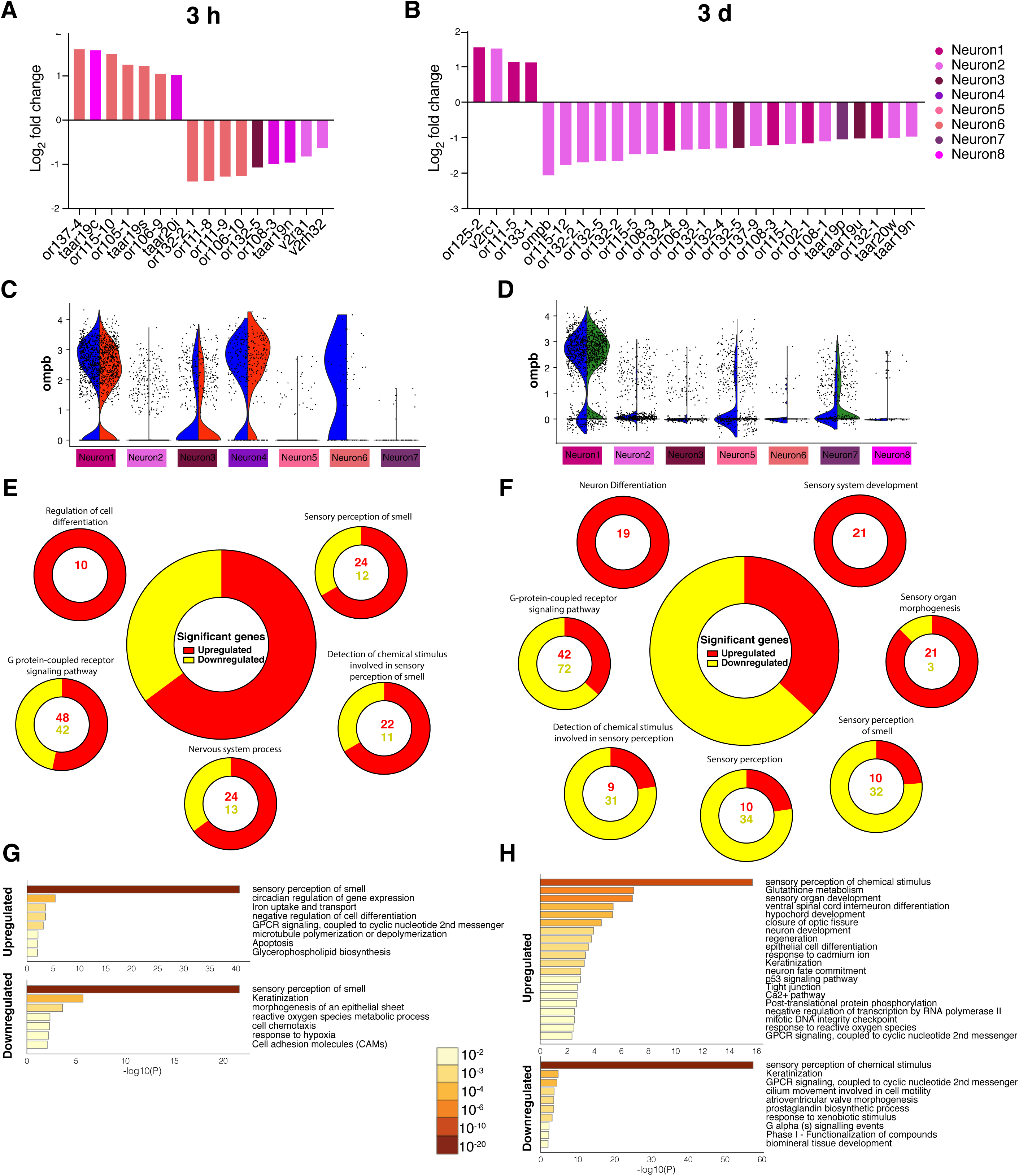
Intranasal SARS-CoV-2 S RBD protein delivery causes widespread losses of olfactory receptor expression in adult zebrafish. (A-B) Bar plots showing fold-change in expression of olfactory receptor genes in the zebrafish olfactory organ 3 h and 3 d post intranasal SARS-CoV-2 S RBD protein compared to vehicle controls identified by scRNA-Seq. (C-D) Violin plots showing changes in *ompb* expression, a marker of mature ciliated OSNs, in zebrafish olfactory organ cell suspensions, across all olfactory neuronal cell clusters in 3 h SARS-CoV-2 S RBD (orange) and 3 d SARS-CoV-2 S RBD (green) versus vehicle treated (blue). (E-F) Summary webs of gene ontology analyses performed in ShinyGO v0.61 from the scRNA-Seq data showing the number of significantly up and down regulated genes across olfactory neuronal clusters in zebrafish olfactory organ 3 h and 3 d post-intranasal SARS-CoV-2 S RBD protein versus vehicle treated controls. (G-H) Gene set enrichment analysis using Metascape of genes from all neuronal clusters that were either significantly up or down regulated in zebrafish olfactory organ 3 h and 3 d post-intranasal SARS-CoV-2 S RBD protein compared to vehicle treated controls (adjusted *P-*value< 0.05). Enriched terms are colored by *P*-value.

### Intranasal delivery of SARS-CoV-2 S RBD induces responses in sustentacular cells and endothelial cell clusters

Co-expression of ACE2 and TMPRSS2 by olfactory SCs and basal cells in the human olfactory epithelium and subsequent onset of damage and inflammation of these cell subsets may explain olfactory loss in COVID-19 patients (Brann et al., 2020; Cooper et al., 2020). Additionally, ACE2 is expressed in ECs and these cells are infected by SARS-CoV-2 (Bryche et al., 2020). Our study allowed us to dissect how each cell type in the zebrafish olfactory organ responds to SARS-CoV-2 S RBD. Our results indicated unique responses by SC clusters and EC clusters to treatment (Figures 4 and 6). At 3 h, we detected increased expression of *apoeb*, of transcription factors *foxq1a* and *id2b*, the transcriptional regulator *nfil3-5*, two tumor necrosis factor receptor superfamily members (*tnfrsf11b* and *tnfrsf9a*) as well as *tcima* (transcriptional and immune response regulator), whose mammalian ortholog *pcim* regulates immune responses as well as endothelial cell activation and expression of inflammatory genes (Figure 6A) (Kim et al., 2009). Further, at 3 h we observed downregulation of the pro-inflammatory cytokine *il17af/3* as well as glutathione peroxidase *gpx1b*, the transcription factor *notch1b*, basal cell adhesion molecule *bcam*, guanine nucleotide-binding protein subunit gamma *gng8*, and the calcium binding *s100z* in SC and EC from SARS-CoV-2 S RBD treated olfactory organs relative to vehicle treated. At 3 d post-treatment, we observed significant increased expression of the gene that encodes brain natriuretic peptide (*nppc*), a vasodilating hormone, the pro-inflammatory chemokine *ccl19a*.*2*, the M2 macrophage marker *arg2*, the transcription factors f*oxq1a* and sox11a, tubulin beta 5 *tubb5*, and the epithelial mitogen *epgn*, among others (Figure 6B). Downregulated genes at 3 d post-treatment included *hsp70*.*3, apoeb*, the osteoblast specific factor b *postb*, the desmosomal component periplakin (*ppl*), the vasoconstricting endothelin 2 (*edn2*), the heparin binding molecule latexin (*ltx*) involved in pain and inflammation, and *cd74b*, a part of the MHC-II complex (Figure 6B). Combined, these data indicated immune regulatory responses in SC and EC clusters early after SARS-CoV-2 S RBD treatment, followed by transcriptional changes with potential vasoactive effects by day 3.

**Figure 6:**
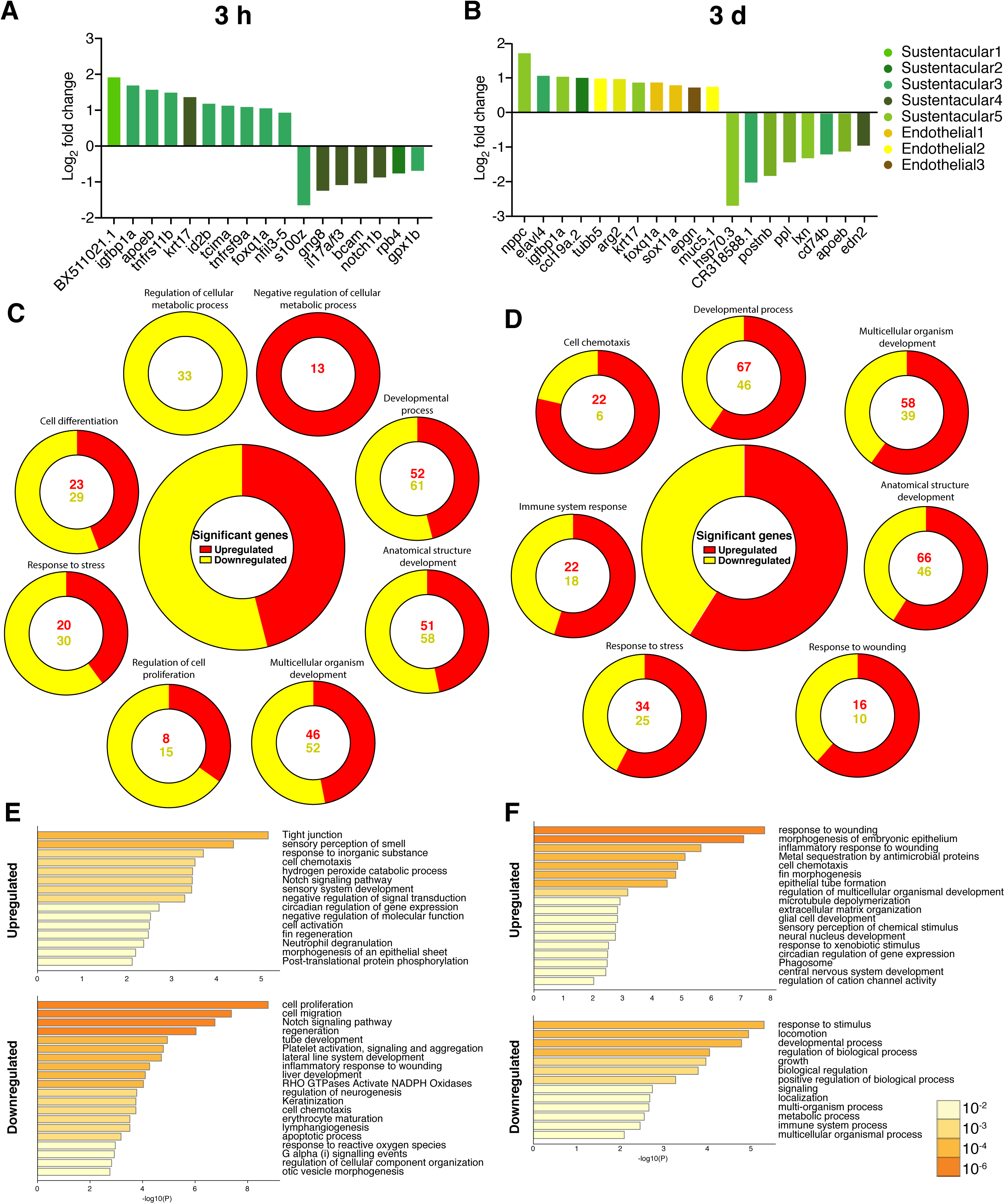
Intranasal SARS-CoV-2 S RBD protein delivery causes transcriptional changes in zebrafish sustentacular and endothelial clusters. (A-B) Bar plots showing fold-change in expression of selected genes in sustentacular and endothelial cell clusters in the zebrafish olfactory organ 3 h and 3 d post intranasal SARS-CoV-2 S RBD protein compared to vehicle controls identified by scRNA-Seq. (C-D) Summary webs of gene ontology analysis performed in ShinyGO v0.61 from the scRNA-Seq data showing the number of genes significantly up and down regulated in sustentacular and endothelial cell clusters in the olfactory organ of zebrafish 3 h and 3 days post-intranasal SARS-CoV-2 S RBD protein compared to vehicle treated controls. (E-F) Gene set enrichment analysis using Metascape of genes from sustentacular and endothelial clusters that were either significantly up or down regulated in zebrafish olfactory organ 3 h and 3 d post-intranasal SARS-CoV-2 S RBD protein compared to vehicle treated controls (adjusted P-value< 0.05). Significantly enriched terms are colored by *P*-value.

GO and enrichment set analyses indicated that SC and EC clusters initially undergo transcriptional changes enriched in metabolic responses, response to stress, and cell differentiation (Figure 6C). Later on, at day 3, SC and EC responses were enriched in genes involved not only in the stress response but also in immune responses and responses to wounding (Figure 6D). Similar results were identified using Metascape, which showed that the inflammatory response to wounding was moderately enriched in the downregulated genes at 3 h, whereas by day 3, response to wounding became the top enriched set among the upregulated genes (Figure 6E-F).

### Exposure of wildtype zebrafish larvae with SARS-CoV-2 does not support viral replication

Zebrafish larvae have been used as models to investigate several human viruses. Infecting zebrafish larvae in a BSL-3 laboratory by immersion in contaminated water is comparable to infecting a cell line. We first checked the stability of SARS-CoV-2 in zebrafish water overtime, in the absence of any animals. We found that SARS-CoV-2 viral loads in the water remained stable throughout the experiment (Figure 7A-B). We exposed wildtype AB zebrafish larvae to live SARS-CoV2 and examined viral mRNA abundance over time to determine if zebrafish larvae can support viral replication. We detected no increases in the viral N copy numbers over time and a steady decline in E gene copy numbers in both water from wells containing larvae and virus as well as in the larval tissue (Figure 7C-F). These results indicate that wild-type zebrafish larvae cannot support efficient SARS-CoV-2 replication as suggested by the *in silico* comparative sequence analyses of the zebrafish ace2 molecule.

**Figure 7:**
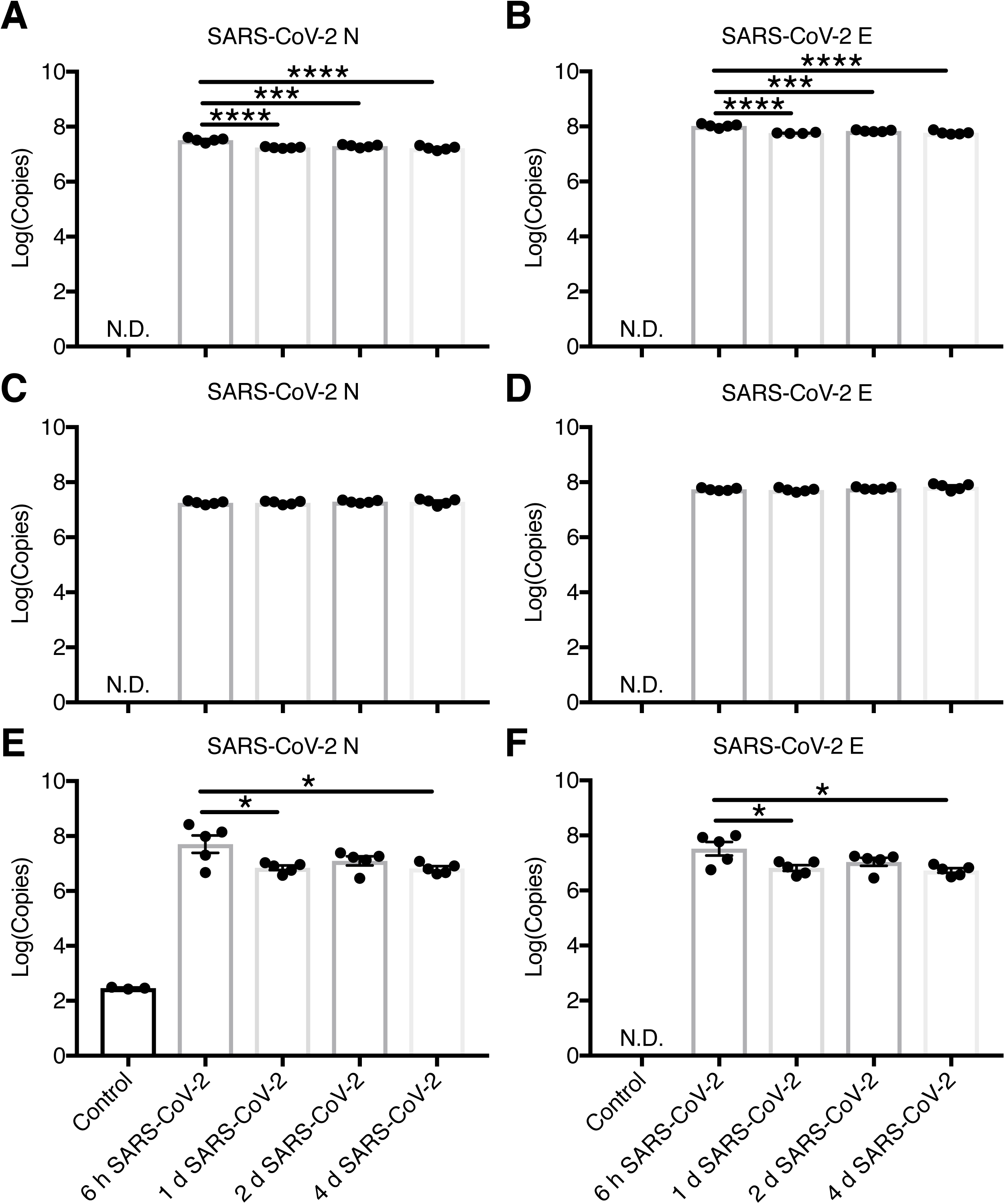
Wildtype zebrafish larvae do not support replication of SARS-CoV-2 virus. (A-B) Mean viral loads quantified as log of SARS-CoV-2 N gene and SARS-CoV-2 E gene copy numbers in control tank water and tank water + 10^4^ pfu of SARS-CoV-2 for 6 h, 1 d, 2 d and 4 d. (C-D) Mean viral loads quantified as log of SARS-CoV-2 N gene and SARS-CoV-2 E gene copy numbers in control supernatants from well with larvae not exposed to virus, and supernatants from wells with larvae that were exposed to 10^4^ pfu of SARS-CoV-2 for 6 h, 1 d, 2 d and 4 d. Each sample represents the supernatant of one well containing 12 larvae. (E-F) Mean viral loads quantified as log of SARS-CoV-2 N gene and SARS-CoV-2 E gene copy numbers in control larvae and larvae exposed to 10^4^ pfu of SARS-CoV-2 for 6 h, 1 d, 2 d and 4 d. Each sample point represents one well containing 12 larvae. Larval infections began at 2 dpf after mechanical dechorionation at 1 dpf. * *P-*value<0.05; ** *P-* value<0.01 *** *P-*value<0.001. Results are representative of two independent experiments.

### Exposure of zebrafish larvae to SARS-CoV-2 decreases ace2 expression and triggers pro-inflammatory cytokine responses

In order to determine whether exposure of zebrafish larvae with live SARS-CoV-2 causes changes in Ai, we measured *ace2* mRNA levels in control and SARS-CoV-2 exposed larvae over time. *ace2* expression was significantly downregulated as early as 6 h post-infection. *ace2* expression inhibition was sustained over the time course of the experiment with the greatest decrease occurring 2 days post-infection (dpi) (Figure 8A). We next evaluated changes in expression of cytokine and chemokine genes to establish whether zebrafish mount inflammatory responses that resemble the patterns of mild or severe SARS-CoV-2 infection. *Il1β* expression was significantly upregulated at 6h, 1 dpi (2-3 fold) and 2 dpi (10 fold) and significantly downregulated at 4 dpi (Figure 8B). We detected a significant increase in *tnfa* expression in SARS-CoV-2 exposed larvae 2 dpi (Figure 8C). *ifnphi1* and *ifnphi3* are the two main type I IFN genes involved in larval zebrafish antiviral responses (Levraud et al., 2019). We detected a significant up-regulation of *ifnphi1* at 1 and 2 dpi, whereas expression was inhibited at 4 dpi. Interestingly, *ifnphi3* expression followed a very different pattern compared to *ifnphi1*, which was significantly downregulated 2 dpi but significantly upregulated at 4 dpi (Figure 8D-E). *Mxa* expression was significantly downregulated at all time points (Figure 8F). *il17af/3* expression was significantly elevated 1, 2 and 4 dpi (Figure 8G). Expression levels of *il22*, a member of the IL10 family, were downregulated 6 hpi and 1 dpi (Figure 8H) whereas the type II cytokine *il4/il13b* was downregulated at 6 hpi, 1 dpi and 2 dpi (Figure 8I). Further, a significant increase in the expression of the chemokine *ccl20a*.*3* was detected in infected larvae at 1 and 2 dpi compared to controls (Figure 8J). A moderate increase in *ccl19a*.*1* expression was observed at 2 dpi followed by a strong down-regulation (15 fold) at 4 dpi (Figure 8K). Taken together, these data indicate that exposure to SARS-CoV-2 induces a significant antiviral and pro-inflammatory immune response in wildtype zebrafish larvae. This response involved type I IFN, *tnfa, il1b, il17* and *ccl20*, reminiscent of COVID-19 patients with mild disease.

**Figure 8:**
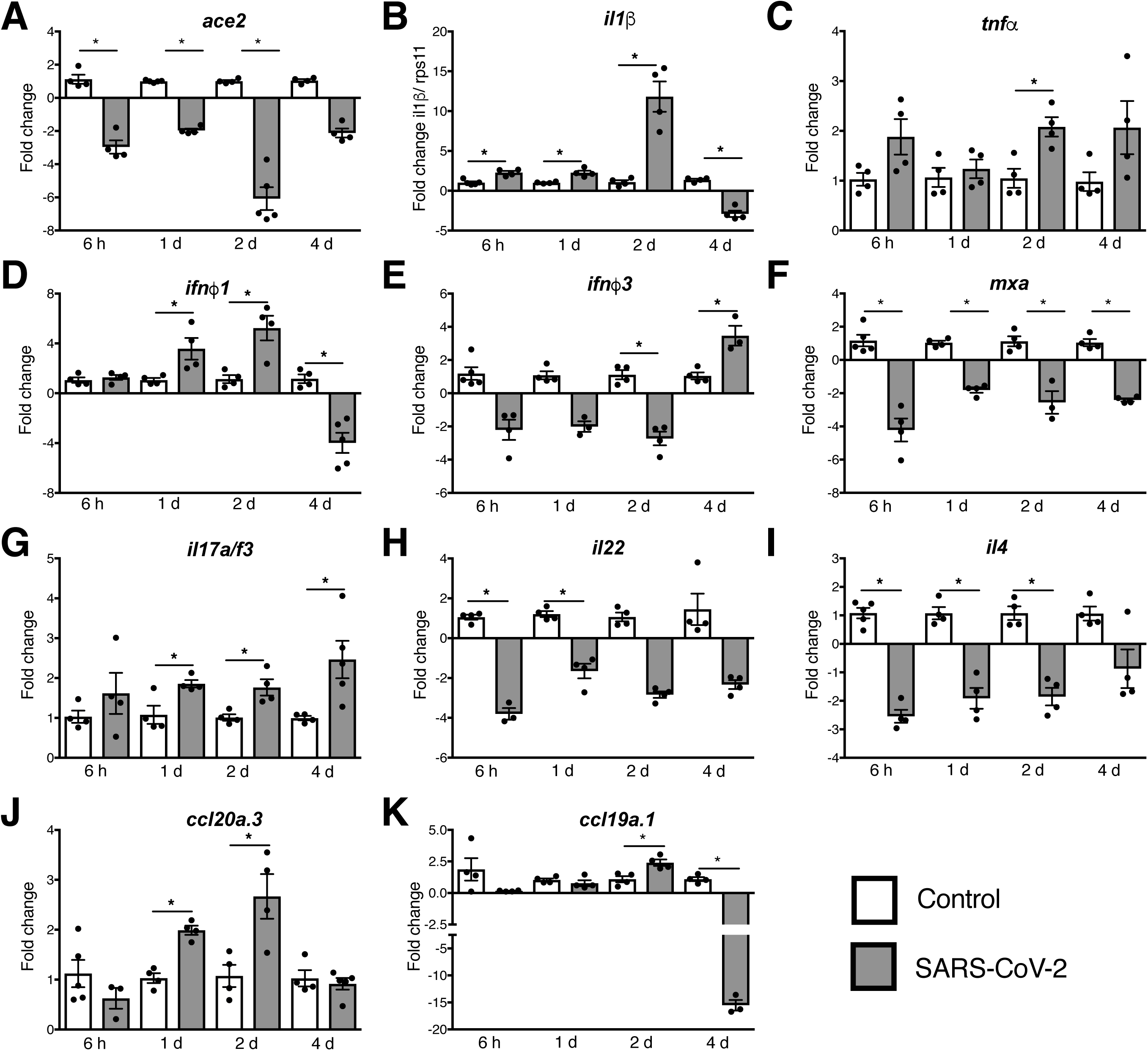
Exposure to SARS-CoV-2 down-regulates *ace2* expression and induces pro-inflammatory immune responses in zebrafish larvae. (A) Changes in *ace2* expression levels in control and SARS-CoV-2 exposed zebrafish. (B) Changes in *il1b* expression in control and infected zebrafish. (C) Changes in *tnfa* expression in control and infected zebrafish. (D) Changes in *ifnphi1* expression in control and infected zebrafish. (E) Changes in *ifnphi3* expression in control and infected zebrafish. (F) Changes in *mxa* expression in control and infected zebrafish. (G) Changes in *il17a/f3* expression in control and infected zebrafish. (H) Changes in *il22*expression in control and infected zebrafish. (I) Changes in *il4/il13* expression in control and infected zebrafish. (J) Changes in *ccl20a*.*3* expression in control and infected zebrafish. (K) Changes in *ccl19a*.*1* expression in control and infected zebrafish. Zebrafish larvae were exposed to 10^4^ pfu of SARS-CoV-2 for 6 h, 1 d, 2 d and 4 d. Larvae incubated at 30□. Infections began at 2 dpf after mechanical dechorionation at 1 dpf. Each sample represents one pool of 12 larvae. Changes in gene expression were calculated as the fold-change compared to vehicle controls using the Pffafl method. Data were normalized using *rps11* expression which was used as the house-keeping gene. * *P-*value<0.05; ** *P-* value<0.01 *** *P-*value<0.001.

## Discussion

The current COVID-19 pandemic has propelled the investigation of SARS-CoV-2 and the development of animal models that help identify therapeutic interventions and vaccines for COVID-19. Thus far, all animal models reported are mammals, and therefore breeding, genetic manipulation, and animal housing in BSL-3 laboratories make these models costly and not readily available in large numbers. Zebrafish can overcome many of the limitations of mammalian models thanks to their transparent bodies, short life-span, low maintenance costs and production of large numbers of embryos. We therefore performed the simplest infection procedure, where SARS-CoV-2 was added to the water of zebrafish larvae. In this manner, BSL-3 trained personnel with no experience in zebrafish microinjection can readily expose larvae to SARS-CoV-2 without the need of animal protocols in a similar fashion to *in vitro* cell culture infections. Exposure of wildtype zebrafish larvae to SARS-CoV-2 in the water did not however result in any detectable viral replication. This finding supports that, despite the general conservation of ACE2 molecules in vertebrates, the amino acid differences identified in zebrafish Ace2, are sufficient to prevent infectivity. Whether infection via microinjection as it was previously done in Influenza A infection models (Gabor et al., 2014) would result in SARS-CoV-2 replication remains to be elucidated. Further, our findings support that, similar to mice, the generation of transgenic larvae expressing human ACE2 may be the best approach in order to support active replication.

Despite the absence of active viral replication, our model provided important insights into the transcriptional innate immune responses to SARS-CoV-2. Pathological cytokine responses are detected in severe disease COVID-19 patients highlighting the need to identify interventions that control such maladaptive inflammatory responses (Blanco-Melo et al., 2020; Hadjadj et al., 2020; Lucas et al., 2020; Del Valle et al., 2020). Mild disease, on the other hand, is characterized by controlled inflammatory and antiviral responses. Zebrafish larvae are known to mount innate immune responses that are largely homologous to those of humans (Zou and Secombes, 2016). In particular, zebrafish possess type I IFNs with similar function and structure to those of mammals and a core ancestral repertoire of IFN-stimulated genes (ISG) has been identified in zebrafish and human (Hamming et al., 2011; Levraud et al., 2019). Zebrafish also possess orthologues to the main inflammatory cytokines (IL1β, TNFα, IL-6, IL-17) and chemokines identified in the response to SARS-CoV-2 in humans (Zou and Secombes, 2016). The present study identified pro-inflammatory cytokine and chemokine signatures in zebrafish larvae exposed to SARS-CoV-2 that largely resemble those described in humans with mild disease symptoms. Interestingly, we detected onset of type 1 pro-inflammatory responses with elevated expression of the cytokines *il1b* and *tnfa*, sustained inhibition of type 2 responses (*il4/il13b*), which would further favor a pro-inflammatory state, and mixed type 3 responses with up-regulation of *il17a/f* but downregulation of *il22*.

Coronaviruses deploy various mechanisms to evade type I IFN responses within infected cells (Park and Iwasaki, 2020; Sa Ribero et al., 2020; De Wit et al., 2016). Interrogation of the main type I IFN genes in zebrafish larvae after co-incubation with live SARS-CoV-2 indicated complex type I IFN responses; significant up-regulation of *ifnphi1* in response to SARS-CoV-2 exposure during the first two days, while ifnphi3 was downregulated. In contrast, at 4dpi, *ifnphi1* was down-regulated but *ifnphi3* expression had increased. Interestingly, *Mxa*, an ISG, was downregulated in response to SARS-CoV-2 exposure. Combined, these data suggest the SARS-CoV-2 induces some type I IFN responses in zebrafish larvae while inhibits others. Future studies are clearly needed to ascertain the role of teleost type I IFN in the anti SARS-CoV-2 immune response.

S protein is a structural protein of SARS-CoV-2 and therefore the target of several vaccine trials. Therefore, we exposed zebrafish larvae to SARS-CoV-2 S RBD protein and investigated transcriptional and physiological responses. Rapid changes in gene expression were detected in treated larvae, including up-regulation of the chemokine c*cl20a*.*3* and the down-regulation of *ifnphi3*. The ccl19/ccl20 axis appears to be critical in teleost antiviral innate responses, as previous studies have shown very rapid responses in larvae exposed to the rhabdovirus SVCV (Sepahi et al., 2019). This change was also detected in the larvae that were exposed to the live SARS-CoV-2 virus in the present study. We further detected a significant down-regulation of type I IFN *ifnphi1* gene in larvae exposed to SARS-CoV-2 S RBD protein.

Examination of *ace2* transcriptional changes in zebrafish larvae exposed to SARS-CoV-2 revealed a consistent down-regulation in expression throughout the course of infection. Interestingly, we did not observe any changes in *ace2* expression after 3h immersion with SARS-CoV-2 S RBD protein. Recently, enterocytes were found to be the main cell type expressing Ace2 in 5 dpf-old zebrafish larvae (Postlethwait et al., 2020); and therefore it is possible that the down-regulation in *ace2* expression observed in our experiments was the result of enterocyte responses to SARS-CoV-2. However, an olfactory epithelial cell cluster was not identified in this dataset, probably because these cells constitute too small a fraction of the cells of an entire larva. Importantly, we exposed larvae to the virus at 3 dpf, when the olfactory pit is already sampling the surrounding water, while the gut fully opens only at 4 dpf. Thus, changes in *ace2* expression levels in the olfactory pit of the zebrafish larvae cannot be ruled out at this point.

Previous work has shown that ACE2 knockdown in mice protects from SARS-CoV infection (Kuba et al., 2005). Thus, down-regulation of zebrafish Ace2 expression may have protected larvae from SARS-CoV-2 infection in our experiments. Our data agree with studies in mouse lungs, where suppression of *ace2* gene expression was consistently observed following SARS-CoV-2 infection (Chen et al., 2020). Interestingly, changes in ACE2 levels can occur in response viruses that do not require ACE2 for host entry (Chen et al., 2020). Thus, although further studies are warranted, our data suggest that Ace2 is involved in antiviral SARS-CoV-2 responses in zebrafish.

We took advantage of the zebrafish fish model which allows for easy live imaging of heart beats in transparent larvae. We detected *in vivo* cardiac/heart responses in larval zebrafish exposed to SARS-CoV-2 S RBD protein characterized by tachycardia. Cardiac arrhythmia is a common symptom among COVID-19 patients and current research efforts aim to understand how SARS-CoV-2 infection impacts cardiovascular function (Libby, 2020). Our findings underscore that SARS-CoV-2 S RBD is able to cause tachycardia in the zebrafish larval model and that this model can be used for rapid evaluation of drug treatments for COVID-19. As a proof of concept, we used captopril, an ACE inhibitor currently being evaluated in human clinical trials (NCT04355429). Captopril treatment ameliorated tachycardia in zebrafish larvae exposed to SARS-CoV-2 S RBD recombinant protein. Our studies therefore suggest the beneficial use of captopril in COVID-19 patients undergoing cardiac arrhythmia, but clearly further studies are required to fully translate these findings to the clinic and to determine the duration and timing of captopril treatment in COVID-19 patients.

A recent report in zebrafish adults indicated that injection of recombinant SARS-CoV-2 S N terminal protein caused histopathology of several tissues including the liver, kidney, brain and ovary (Ventura-Fernandes et al., 2020). Additionally, some animals succumbed to injection with the recombinant protein. We did not detect any mortalities neither in larvae nor in adults in any of our experiments, perhaps suggesting that mortalities were due to the injection procedure rather than the protein treatment itself. Of note, the dose used in the present study was considerably lower than the dose delivered in the Ventura-Fernadez study, perhaps explaining the differences in toxicity between both studies. We observed histological damage following a single intranasal delivery of SARS-CoV-2 protein, specifically at the site of delivery, the olfactory organ, whereas more transient and moderate damage was detected in the renal tubules. Renal damage may have occurred by direct uptake of SARS-CoV-2 S RBD by the kidney once the protein reached the bloodstream following intranasal administration or, alternatively, by activation of RAS or inflammatory cascades at the olfactory organ. Thus, toxicity of SARS-CoV-2 S protein in adult zebrafish may be less severe when delivered intranasally than by injection, and future studies should evaluate whether current vaccine candidates also exert similar effects and whether different administration routes cause the same side-effects or not. This is particularly important as the intranasal route appears promising for some vaccine candidates (Ku et al., 2020).

SARS-CoV-2 infection causes anosmia and ageusia in 41-64% of patients (Dell’Era et al., 2020; Spinato et al., 2020). Several mechanisms for the impaired chemosensory perception in COVID-19 patients have been proposed based on the observation that, in humans, sustentacular cells and basal cells but not OSNs co-express *ace2* and *tmprss2* (Brann et al., 2020). Initial infection of SCs may indirectly cause damage to OSNs, notably via lack of metabolic support or via inflammatory-derived damage (Brann et al., 2020). Animal models of SARS-CoV-2 infection have provided some insights into the mechanisms of olfactory loss driven by SARS-CoV-2. In golden Syrian hamster, intranasal SARS-CoV-2 infection results in drastic damage of the olfactory epithelium including ciliary loss. In this animal model, while a study reported infection of sustentacular cells but not OSNs, another study claimed direct infection of both mature and immature OSNs (Zhang et al., 2020). In the Syrian hamster model, studies found infiltration of inflammatory cells into the olfactory lamina propria (Bryche et al., 2020). Our results indicate that intranasal delivery of SARS-CoV-2 S RBD is sufficient to cause severe olfactory damage including loss of cilia, edema, hemorrhages and necrosis in adult zebrafish. However, no inflammatory infiltrates were observed in our model compared to the hamster model, likely due to the absence of an active viral infection. scRNA-Seq of the zebrafish olfactory organ identified dramatic losses in OR expression in most OSN subsets. As a result, intranasal delivery of SARS-CoV-2 S RBD caused changes in OSN populations characterized by loss of ciliated and microvillous mature OSNs and increased proportions of cycling immature OSNs 3 d post-treatment. Further, inflammatory gene signatures could be detected in SCs, ECs and myeloid cell subsets in animals treated with SARS-CoV-2 S RBD. The broad loss in OR, VR and TAAR expression in the zebrafish olfactory organ was paralleled by sudden hyposmia and anosmia according to functional EOG recordings. This finding largely recapitulates previous results in K18-hACE2 mice, which develop anosmia soon after infection (Zhen et al., 2020). The present study did not determine when zebrafish recover olfactory function following SARS-CoV-2 S RBD intranasal treatment, but based on our histopathological observations, the olfactory organ was still severely damaged 5 days after treatment, suggesting that recovery of olfactory function may take several weeks in our model. Our findings therefore indicate that similar to humans, zebrafish suffer from olfactory pathology and loss of smell in response to SARS-CoV-2 S RBD protein. Thus, olfactory pathophysiology appears to occur even in the absence of viral replication raising the possibility that nasal vaccines for COVID-19 may also cause transient anosmia in humans.

In conclusion, the present study reports that both adult and larval wild-type zebrafish can be useful models to advance our understanding of COVID-19 pathophysiology, particularly of mild disease. The present study is the foundation for the use of zebrafish as a novel vertebrate model that can accelerate the investigation of SARS-CoV-2 pathophysiology, decipher the dynamics of host innate immune responses and test drugs in a fast, inexpensive and high-throughput manner. Identification of molecules with antiviral effects against SARS-CoV-2 is typically done using *in vitro* cell cultures (Wang et al., 2020b). However, such *in vitro* approaches do not provide organ specific information or any physiologically relevant readouts. Thus, the zebrafish model here established can serve as a platform to screen and develop drug interventions for COVID-19 where whole-organismal *in vivo* readouts can be generated in very short periods of time and in an inexpensive manner. Further, zebrafish could be used in high throughput screens in which not only drugs but also genetic and environmental risk factors known to impact COVID-19 severity can be incorporated during drug validation and testing.

## Acknowledgments

Authors thank Dr. F. Krammer for kindly sharing the SARS-CoV-2 spike RBD recombinant protein. Thank you to Dr. Ryan Wong for providing some of the AB adult animals used in this study. This study was funded by NSF award # #1755348.

## Author Contributions

Conceptualization, A.K., I.S., P.B., and JP.L., Methodology, A.K., I.S., M.H., S.B, JP.L.; Investigation, A.K., I.S., M.H., S.B, JP.L.; Writing –Original Draft, A.K and I.S.; Writing – Review & Editing, A.K., I.S., M.H.,W.L., S.B., P.B., and JP.L.; Funding Acquisition, I.S., M.H., and S.B.; Resources, I.S., M.H., S.B. JP.L.; Supervision, I.S.

## Declaration of Interests

Authors declare no competing interests

## EXPERIMENTAL MODEL AND SUBJECT DETAILS

### Ethics statement

All animal protocols were done in accordance with American Veterinary Medical Association guidelines under a protocol approved by the University of New Mexico’s Institutional Animal Care and Use Committee (Protocol Number 19-200863-MC), the University of New Mexico Health Sciences Center Institutional Biosafety Committee (Protocol Number 17-11-01R) and the Texas State University Institutional Animal Care and Use Committee (Protocol Number 43).

### Zebrafish Husbandry

For all experiments, wild type AB zebrafish were obtained from ZIRC (Oregon, USA). For the intranasal delivery of SARS-CoV-2 S RBD protein into adult zebrafish, female and male adult zebrafish were obtained from Dr. Wong’s laboratory at the University of Nebraska due to lockdown of ZIRC during the pandemic. All fish were maintained in a filtered aquarium system at 28°C with a 14 h light and 10 h dark cycle at the University of New Mexico Aquatics Animal Facility. All experiments with adults utilized a mix of male and female animals, and the larvae sex is indeterminable. Animals were fed *ad libitum* Gemma complete nutrition (Skretting). AB larvae were obtained by batch-crossing AB adults allowing for natural fertilization. The morning of fertilization, larvae were collected at n=50 per Petri dish and kept in E3 medium containing 0.002% methylene blue. In the afternoon, larvae were placed in fresh E3 medium without methylene blue and non-surviving embryos were removed. Larvae were maintained at 28.5°C in E3 medium until 4dpf when they are slowly changed to system water.

### SARS-CoV-2

The SARS-CoV-2 isolate, a CDC isolate from a US patient (USA-WA1/2020), was obtained from BEI Resources. The strain was grown at a low MOI to minimize generation of noninfectious particles and low passaged virus was used in all experiments described. The virus was propagated on Vero E6 cells and viral loads quantitated by RT-qPCR and by plaque forming assays as we have described previously (Bradfute et al, 2020).

## METHOD DETAILS

### Intranasal delivery of SARS-CoV-2 S RBD recombinant protein to adult zebrafish

Adult zebrafish were anesthetized for 1 min in 0.04 mg/ml Tricaine-S (Syndel) solution and then moved to an absorbent boat where their gills were still covered with anesthetic solution for administration of solutions to nares. Using a microloader tip (Eppendorf, 930001007), 5 μl of 20 ng/μl SARS-CoV-2 S RBD (kindly provided by Dr. F. Krammer) was directly pipetted into each naris, while 5 μl of sterile PBS was applied in control fish. After inoculation, animals were recovered in a separate tank supplemented with O2 before returning to their rearing tank until the end of the experiment. Euthanasia was performed on ice to ensure rapid death without perturbing the olfactory system.

### Histology and image analysis

Adult zebrafish heads (n=3) were fixed in 4% formaldehyde solution buffered to pH 7.4 with 60 mM HEPES overnight at 4°C on a low speed rocker. Heads were then rinsed three times with PBS for 30 min at room temperature (RT) on a slow rocker and decalcified in 10% EDTA solution (pH 7.4) at RT for 24 h. After rinsing in PBS, samples were placed in 70% ethanol for 24 h at 4°C and then processed for embedding in paraffin according to ZIRC protocols. Paraffin blocks were sectioned at 5-μm thickness and stained by hematoxylin and eosin (H&E) staining. Sections were observed under a Nikon Ti inverted microscope (brightfield), and images were acquired with the Nikon Elements Advanced Research software and analyzed using ImageJ.

### Isolation of olfactory organ cells from adult zebrafish, scRNA-Seq, scRNA-Seq data analysis and Gene Ontology

To obtain single cell suspensions, adult AB zebrafish olfactory rosettes were dissected and placed directly into 5 ml of Neurobasal medium (Gibco, 21103049) supplemented with 10% heat inactivated fetal bovine serum (FBS, Peak Scientific, Colorado, USA), 100 U/ml penicillin-streptomycin (Gibco, 15140122), and 0.04% filtered bovine serum albumin (Sigma) and rocked at 4°C. Every 20 min, the supernatant from each sample was collected and replaced by fresh medium for a total of 4 times. Supernatants were then filtered through a cell strained and kept on ice. Next, the rosettes were placed in 10 ml of a DTT solution (1.3 mM EDTA, 864 μM DTT in PBS) at RT for 30 min. Suspensions were filtered through a cell strainer and combined with the supernatants obtained from the first steps. Rosettes were then rinsed off the DTT solution and incubated in Hank’s buffer containing 10% FBS, 0.04% BSA, 100U/ml penicillin-streptomycin and collagenase type IV (Gibco, 0.37 mg/ml) for 1.5 h at RT. The supernatants were filtered through a cell strainer and combined with those obtained from the previous steps while the rosettes were mashed against the cell strainer and washed with neurobasal medium. The combined supernatants were centrifuged for 10 min at 400g in supplemented neurobasal medium and cells were counted with a hemocytometer. Viability was estimated by trypan blue staining. Cells were then strained twice through Flowmi 10 μm strainers and loaded onto the chromium controller with a viability of > 85%. Cell libraries were generated according to 10x Genomics protocols at the University of New Mexico Cancer Center Genomics core facility and sequenced on an Illumina NovaSeq 6000 at the University of Colorado Genomics and Microarray core facility. Sequencing depth and statistics of the scRNA-Seq run are shown in Figure S2. SRAs for this project can be found on NCBI under bioproject #PRJNA668529.

Fastqs were run through the Cell Ranger v3.0 pipeline with default settings using the GRCz11 zebrafish genome. Output matrices were loaded to R (v1.2.5001) as a Seurat object (Package Seurat v3.1.1). First, cells with less than 200 or greater than 2500 features, and greater than 5% mitochondrial features were removed. After counts were normalized using the “lognormalize” method and a scale factor of 10000, 2000 variable genes were selected using the ‘vst” method. Data was scaled, and PCA dimensional reduction was run. Jackstraw analysis determined the vehicle control to have 38 significant principal components (PCs) and the treated samples to have 40 significant PCs which were used for clustering analysis. SARS-CoV-2 S RBD treated samples were integrated with the vehicle treated sample and clustered together using 30 significant PCs and a resolution of 0.5. Cluster markers were identified with “FindAllMarkers” in Seurat and exported for cluster identification. Differential expression analysis was done with seurat “FindMarkers” in default settings for each cluster and exported for gene ontology analysis.

Gene ontology (GO) analysis was done with web-based GUIs Metascape and ShinyGO v0.61 which draw multiple currently maintained databases (ensembl, ENTRZ, KEGG among others). Biological process webs were created using biological process output from ShinyGO v0.61 in Prism GraphPad. Biological processes bar graphs were produced by Metascape.

### Electro-olfactogram recordings

Adult AB zebrafish were anesthetized and received 2 µl of recombinant SARS-CoV-2 S RBD protein (50 ng/µl) in PBS or PBS alone. After 3 h or 1 day, zebrafish were anaesthetized (0.1 g MS222/L), placed in a v-shape stand and supplied with aerated water containing MS222 anesthetic (0.05 g/L). The nasal flap was removed with sterile fine forceps to expose the olfactory rosettes to a continuous tank water source. Olfactory responses to zebrafish food extract or goldfish bile were measured by electrical recordings as detailed in Sepahi et al., 2019. The food extract was prepared as a filtered solution of 1L tap water and 0.1 g of dry food pellets. Water food extracts were separated in 200 ml aliquots and kept frozen until the recording day. A 0.5 ml mix of bile fluid from 50 adult goldfish was aliquoted in 10 µl and kept frozen until the day of the recording. Before each recording, bile aliquots were diluted 1:1000 in water from the EOG system. There were no significant differences in olfactory responses between males and females, hence responses of both sexes were averaged together. The percentage reduction in olfactory activity was calculated by dividing the amplitude of the olfactory signal at time x by amplitude of the olfactory signal at time 0 *100. Percentage of olfactory signal reduction between control and treated naris was calculated as follows (Amplitude response to odorant in control naris (mV) - Amplitude response to odorant in treated naris (mV))/Amplitude response to odorant in control naris (mV)) * 100.

### Exposure of zebrafish larvae to SARS-CoV-2 S RBB recombinant protein and captopril treatment

5 dpf or 7 dpf zebrafish (n=5/well, total of n=7-8 replicate wells) were placed in a 12 flat-bottomed well plate (FALCON) with 2 ml of tank water per well. Wells were then treated with either 2 ng/ml of recombinant SARS-CoV-2 S RBD, 2 ng/ml recombinant IHNVg protein (Sepahi et al., 2019), with or without 12 µM captopril (Sigma-Aldrich, C4042). The corresponding vehicle controls consisted of sterile PBS for recombinant proteins and water for captopril. Animals were incubated at 28.5°C for 3 h, mounted in 1% low melting point agarose (Sigma) at 42°C, stabilized for 5 min at RT and then imaged to record heart-beat activity.

### Zebrafish larvae heart-beat recordings and analysis

As the agarose solidified, animals were adjusted to the microscope stage (approx. 5 min) then hearts were recorded using brightfield AVI for 3 min at 16.667 frames/s at RT. AVI images were then opened in ImageJ with the Time series analyzer V3 plugin. Circular ROIs were drawn in either the atrium or ventricle and average intensity was extracted. The maximum average intensity peaks were identified and counted per 60s as bpm. Data were analyzed by one-way ANOVA with Tukey’s post hoc.

### Infection of zebrafish fish larvae with SARS-CoV-2

Ten animals were placed in each well in 12-well plates containing 2 ml of tank water and transferred to BSL3 facility the day before infection. Infection was performed by adding with 10^4^ plaque forming units (pfu)/well in a volume of approximately 2 µl. Larvae were collected 6 hpi, 1 dpi, 2 dpi, or 4 dpi in Trizol for RNA extraction. Water (supernatants) from all wells was collected in AVL buffer (Qiaamp viral RNA mini kit, Qiagen) with carrier RNA at 6 hpi, 1 dpi, 2 dpi, or 4 dpi.

### Quantification of SARS-CoV-2 viral loads

RNA from tank water was extracted using the Qiaamp viral RNA mini kit (Qiagen, 52904) according to manufacturer’s instruction, with one modification: after the AVL buffer was frozen and thawed, the solution was the heated to 50°C for 5 min and allowed to come to RT before use. RT-qPCR of the N and E genes of SARS-CoV-2 was done using research use only primer probes sets from IDT (10006713 (N) and 10006888, 10006890, and 10006892 (E)) with plasmid positive controls for a standard curve (10006625 and 10006896, respectively). 5 µl of RNA eluted from the Qiaamp columns was added for each working reaction with Taqman Fast Virus one-step master mix (4444432) and appropriate working concentrations of each primer/probe set (N: 500 nM primers and 125 nM probe; E: 400 nM primers and 200 nM probe). Plates were run on BioRad C1000 touch and CEX real time systems according to Taqman master mix’s protocol and analyzed using Biorad CFX qPCR software.

### Gene expression analyses by RT-qPCR

Whole larvae RNA was extracted using trizol. For tissue homogenization bead beater tubes are preloaded with 1.5 g 1.0 mm dia Zirconia beads, 1.5 g 2.0 mm Zirconia beads and 500 µl Trizol. Samples were loaded into the tubes and bead beat at 4350 rpm for 45 s. Tubes were then centrifuged at 7,000 rpm for 7 min. The homogenate/lysates were transferred to clean 1.5 ml microfuge tubes and spun at 7,000 rpm for 7 min to pellet debris. Supernatants were then processed to extract the total RNA using a standard chloroform/phenol extraction protocol. RNA was quantified by Nanodrop and samples were normalized and 1 µg of RNA was used to synthesize cDNA using the Superscript III first strand system (Thermofisher, 18080051). qPCR was performed using SSOadvanced supermix (Biorad, 1725270) and primers listed in Table S3 (Supplemental Methods). Gene expression changes were quantified using the Pfaffl method (Pfaffl, 2001).

## QUANTIFICATION AND STATISTICAL ANALYSIS

### Statistical analysis

Statistics were done in GraphPad Prism V7.0d. Unpaired t-tests or one-way ANOVA followed by a Tukey’s post hoc comparison was used to detect differences among treatments. All data were checked for normal distribution prior to performing statistical analyses using F test or Bartlett’s. The *P-*values are as follows: * *P-*value < 0.05, ** *P-*value < 0.01, *** *P-*value < 0.001, and **** *P-*value < 0.0001.

### Supplemental Information titles and legends

Table S1: Phylogenetic sequence analyses of vertebrate ACE2 amino acid sequences showing the percent similarity and homology. Analyses were performed in MATGAT. *Homo sapiens* (human), *Macaca mulatta* (macaque), *Mus musculus* (mouse), *Rattus norvegicus* (rat), *Mustela putorius furo* (ferret), *Manis javanica* (pangolin), *Rhinolophus sinicus* (bat), *Camelus dromedaries* (camel), *Canis familiaris* (dog), *Gallus gallus* (chicken), *Alligator mississippiensis* (alligator), *Xenopus tropicalis* (frog), *Danio rerio* (zebrafish).

Table S2: Sequence analyses of SARS-CoV-2 S protein-ACE2 binding amino acids in different vertebrates. Amino acids in red denote non-conserved amino acids. In green, amino acid substitutions that represent non-functional changes.

Table S3: List of primers used in the present study.

Figure S1: Intranasal delivery of SARS-CoV-2 S RDB protein causes renal injury in adult zebrafish. H&E stains of zebrafish head-kidney paraffin sections. (A) Control vehicle treated head-kidney. (B) 3 h; (C) 1 day; (D) 5 days post intranasal delivery of SARS-CoV-2 S RDB protein. Arrows indicate vacuolization of tubule epithelial cells. Images representative n=10 sections per animal and n=3 per experimental group. PT: proximal tubule. DT: distal tubule.

Figure S2: scRNA-Seq identification of cell clusters in zebrafish olfactory organ based on previously established homologous markers.

(A) Table of metrics from the scRNA-Seq. Cells were captured with 10x chromium platform and libraries sequenced on an Illumina NovaSeq 6000.

(B) Heatmap showing markers for each of the 20 cell clusters present in the olfactory organ of adult zebrafish following vehicle treatment and 3 h post intranasal delivery of SARS-CoV-2 S RDB.

(C) Heatmap of markers for each of the 22 cell clusters present in the olfactory organ of adult zebrafish following vehicle treatment and 3 d post intranasal delivery of SARS-CoV-2 S RDB.

Figure S3: Basal expression levels of *ace2* in wildtype adult zebrafish olfactory organ (OO) and olfactory bulb (OB) (n=3) quantified by RT-qPCR. Data were normalized to *rps11* as the house keeping gene.

## Notes

### Competing Interest Statement

The authors have declared no competing interest.

## References

Advani, A. (2020). Acute Kidney Injury: A Bona Fide Complication of Diabetes. Diabetes 69 (11), 2229–2237.

Albini, A., Di Guardo, G., Noonan, D.M.C., and Lombardo, M. (2020). The SARS-CoV-2 receptor, ACE-2, is expressed on many different cell types: implications for ACE-inhibitor- and angiotensin II receptor blocker-based cardiovascular therapies. Intern. Emerg. Med. 15, 759–766.

Babapoor-farrokhran, S., Gill, D., Walker, J., Rasekhi, R.T., Bozorgniaa, B., and Amanullaha, A. (2020). Myocardial injury and COVID-19: Possible mechanisms Savalan. Life Sci. 253, 117723.

Bao, L., Deng, W., Huang, B., Gao, H., Liu, J., Ren, L., Wei, Q., Yu, P., Xu, Y., Qi, F., et al. (2020). The pathogenicity of SARS-CoV-2 in hACE2 transgenic mice. Nature 583, 830–833.

Bergsland, M., Werme, M., Malewicz, M., Perlmann, T., and Muhr, J. (2006). The establishment of neuronal properties is controlled by Sox4 and Sox11. Genes Dev. 20, 3475–3486.

Bermingham-McDonogh, O., and Reh, T.A. (2011). Regulated reprogramming in the regeneration of sensory receptor cells. Neuron 71, 389–405.

Blanco-Melo, D., Nilsson-Payant, B.E., Liu, W.-C., Uhl, S., Hoagland, D., Møller, R., Jordan, T.X., Oishi, K., Panis, M., Sachs, D., et al. (2020). Imbalanced Host Response to SARS-CoV-2 Drives Development of COVID-19. Cell 181, 1036–1045.

Borders, A.S., Hersh, M.A., Getchell, M.L., Van Rooijen, N., Cohen, D.A., Stromberg, A.J., and Getchell, T. V. (2007). Macrophage-mediated neuroprotection and neurogenesis in the olfactory epithelium. Physiol. Genomics 31, 531–543.

Brann, D.H., Tsukahara, T., Weinreb, C., Lipovsek, M., Van Den Berge, K., Gong, B., Chance, R., Macaulay, I.C., Chou, H.J., Fletcher, R.B., et al. (2020). Non-neuronal expression of SARS-CoV-2 entry genes in the olfactory system suggests mechanisms underlying COVID-19- associated anosmia. Sci. Adv. 6, 1–20.

Bryche, B., St Albin, A., Murri, S., Lacôte, S., Pulido, C., Ar Gouilh, M., Lesellier, S., Servat, A., Wasniewski, M., Picard-Meyer, E., et al. (2020). Massive transient damage of the olfactory epithelium associated with infection of sustentacular cells by SARS-CoV-2 in golden Syrian hamsters. Brain. Behav. Immun. 89, 579–586.

Buiakova, O.I., Baker, H., Scott, J.W., Farbman, A., Kream, R., Grillo, M., Franzen, L., Richman, M., Davis, L.M., Abbondanzo, S., et al. (1996). Olfactory marker protein (OMP) gene deletion causes altered physiological activity of olfactory sensory neurons. Proc. Natl. Acad. Sci. U. S. A. 93, 9858–9863.

Burgos, J.S., Ripoll-Gomez, J., Alfaro, J.M., Sastre, I., and Fernando Valdivieso (2008). Zebrafish as a New Model for Herpes Simplex Virus Type 1 Infection. Zebrafish 5, 323–333.

Butler, A., Hoffman, P., Smibert, P., Papalexi, E., and Satija, R. (2018). Integrating single-cell transcriptomic data across different conditions, technologies, and species. Nat. Biotechnol. 36, 411–420.

Chan, J.F.W., Zhang, A.J., Yuan, S., Poon, V.K.M., Chan, C.C.S., Lee, A.C.Y., Chan, W.M., Fan, Z., Tsoi, H.W., Wen, L., et al. (2020). Simulation of the clinical and pathological manifestations of Coronavirus Disease 2019 (COVID-19) in golden Syrian hamster model: implications for disease pathogenesis and transmissibility. Clin. Infect. Dis. 2019. ciaa325.

Chen, J., Jiang, Q., Xia, X., Liu, K., Yu, Z., Tao, W., Gong, W., and Han, J.D.J. (2020). Individual variation of the SARS-CoV-2 receptor ACE2 gene expression and regulation. Aging Cell 19, 1–12.

Chen, M., Reed, R.R., and Lane, A.P. (2017). Acute inflammation regulates neuroregeneration through the NF-κB pathway in olfactory epithelium. Proc. Natl. Acad. Sci. 114, 8089–8094.

Chen, M., Reed, R.R., and Lane, A.P. (2019). Chronic Inflammation Directs an Olfactory Stem Cell Functional Switch from Neuroregeneration to Immune Defense. Cell Stem Cell 25, 501–513.e5.

Cooper, K.W., Brann, D.H., Farruggia, M.C., Bhutani, S., Pellegrino, R., Tsukahara, T., Weinreb, C., Joseph, P. V., Larson, E.D., Parma, V., et al. (2020). COVID-19 and the Chemical Senses: Supporting Players Take Center Stage Keiland. Neuron 107, 219–233.

Damas, J., Hughes, G.M., Keough, K.C., Painter, C.A., Persky, N.S., Corbo, M., Hiller, M., Koepfli, K.P., Pfenning, A.R., Zhao, H., et al. (2020). Broad host range of SARS-CoV-2 predicted by comparative and structural analysis of ACE2 in vertebrates. Proc. Natl. Acad. Sci. U. S. A. 117, 22311–22322.

Dell’Era, V., Farri, F., Garzaro, G., Gatto, M., Aluffi Valletti, P., and Garzaro, M. (2020). Smell and taste disorders during COVID-19 outbreak: Cross-sectional study on 355 patients. Head Neck 42, 1591–1596.

Del Valle, D.M., Kim-Schulze, S., Huang, H.H., Beckmann, N.D., Nirenberg, S., Wang, B., Lavin, Y., Swartz, T.H., Madduri, D., Stock, A., et al. (2020). An inflammatory cytokine signature predicts COVID-19 severity and survival. Nat. Med. 26, 1636–1643.

Ellett, F., Pase, L., Hayman, J.W., Andrianopoulos, A., and Lieschke, G.J. (2011). Mpeg1 Promoter Transgenes Direct Macrophage-Lineage Expression in Zebrafish. Blood 117, 49–56.

Fu, J., Zhou, B., Zhang, L., Balaji, K.S., Wei, C., Liu, X., Chen, H., Peng, J., and Fu, J. (2020). Expressions and significances of the angiotensin-converting enzyme 2 gene, the receptor of SARS-CoV-2 for COVID-19. Mol. Biol. Rep. 47, 4383–4392.

Gabor, K.A., Goody, M.F., Witten, P.E., Mowel, W.K., Breitbach, M.E., Kim, C.H., and Gratacap, R.L. (2014). Influenza A virus infection in zebrafish recapitulates mammalian infection and sensitivity to anti-influenza drug treatment. Dis. Model. Mech. 7, 1227–1237.

Galindo-Villegas, J. (2020). COVID-19: Real-time dissemination of scientific information to fight a public health emergency of international concern. Front. Pharmacol. 11, 10–11.

Garvin, M.R., Alvarez, C., Miller, J.I., Prates, E.T., Walker, A.M., Amos, B.K., Mast, A.E., Justice, A., Aronow, B., and Jacobson, D. (2020). A mechanistic model and therapeutic interventions for COVID-19 involving a RAS-mediated bradykinin storm. eLife 2020; 9:e59177.

Ge, S.X., Jung, D., and Yao, R. (2020). ShinyGO: a graphical gene-set enrichment tool for animals and plants. Bioinformatics 36, 2628–2629.

Giancotti, V., Bergamin, N., Cataldi, P., and Rizzi, C. (2018). Epigenetic Contribution of High-Mobility Group A Proteins to Stem Cell Properties. Int. J. Cell Biol. 2018, 3698078.

Glowacka, I., Bertram, S., Herzog, P., Pfefferle, S., Steffen, I., Muench, M.O., Simmons, G., Hofmann, H., Kuri, T., Weber, F., et al. (2010). Differential Downregulation of ACE2 by the Spike Proteins of Severe Acute Respiratory Syndrome Coronavirus and Human Coronavirus NL63. J. Virol. 84, 1198–1205.

Gomes, M.C., and Mostowy, S. (2020). The Case for Modeling Human Infection in Zebrafish. Trends Microbiol. 28, 10–18.

Guzik, T.J., Mohiddin, S.A., Dimarco, A., Patel, V., Savvatis, K., Marelli-Berg, F.M., Madhur, M.S., Tomaszewski, M., Maffia, P., D’Acquisto, F., et al. (2020). COVID-19 and the cardiovascular system: Implications for risk assessment, diagnosis, and treatment options. Cardiovasc. Res. 116, 1666–1687.

Hadjadj, J., Yatim, N., Barnabei, L., Corneau, A., Boussier, J., Pere, H., Charbit, B., Bondet, V., Chenevier-Gobeaux, C., Breillat, P., et al. (2020). Impaired type I interferon activity and exacerbated inflammatory responses in severe Covid-19 patients. Science 369, 718–724.

Hamming, O.J., Lutfalla, G., Levraud, J.-P., and Hartmann, R. (2011). Crystal Structure of Zebrafish Interferons I and II Reveals Conservation of Type I Interferon Structure in Vertebrates. J. Virol. 85, 8181–8187.

Hansen, A., Rolen, S.H., Anderson, K., Morita, Y., Caprio, J., and Finger, T.E. (2003). Correlation between Olfactory Receptor Cell Type and Function in the Channel Catfish. J. Neurosci. 23, 9328–9339.

Hoffmann, M., Kleine-Weber, H., and Pöhlmann, S. (2020a). A Multibasic Cleavage Site in the Spike Protein of SARS-CoV-2 Is Essential for Infection of Human Lung Cells. Mol. Cell 78, 779–784.e5.

Hoffmann, M., Kleine-Weber, H., Schroeder, S., Krüger, N., Herrler, T., Erichsen, S., Schiergens, T.S., Herrler, G., Wu, N.H., Nitsche, A., et al. (2020b). SARS-CoV-2 Cell Entry Depends on ACE2 and TMPRSS2 and Is Blocked by a Clinically Proven Protease Inhibitor. Cell 181, 271–280.e8.

Howe, K., Clark, M.D., Torroja, C.F., Torrance, J., Berthelot, C., Muffato, M., Collins, J.E., Humphray, S., McLaren, K., Matthews, L., et al. (2013). The zebrafish reference genome sequence and its relationship to the human genome. Nature 496, 498–503.

Imai, Y., Kuba, K., Rao, S., Huan, Y., Guo, F., Guan, B., Yang, P., Sarao, R., Wada, T., Leong-Poi, H., et al. (2005). Angiotensin-converting enzyme 2 protects from severe acute lung failure. Nature 436, 112–116.

Imam, S.A., Lao, W.P., Reddy, P., Nguyen, S.A., Schlosser, R.J. (2020). Is SARS-CoV-2 (COVID-19) postviral olfactory dysfunction (PVOD) different from other PVOD? World J. Otorhinolaryngol. Head Neck Surg. 2020;10.1016/j.wjorl.2020.05.004. doi:10.1016/j.wjorl.2020.05.004

Jia, H., Yue, X., and Lazartigues, E. (2020). ACE2 mouse models: a toolbox for cardiovascular and pulmonary research. Nat. Commun. 11, 5165.

Kim, J., Kim, Y., Kim, H.-T., Kim, D.W., Ha, Y., Kim, J., Kim, C.-H., Lee, I., and Song, K. (2009). TC1(C8orf4) Is a Novel Endothelial Inflammatory Regulator Enhancing NF-κB Activity. J. Immunol. 183, 3996–4002.

Kim, Y. Il, Kim, S.G., Kim, S.M., Kim, E.H., Park, S.J., Yu, K.M., Chang, J.H., Kim, E.J., Lee, S., Casel, M.A.B., et al. (2020). Infection and Rapid Transmission of SARS-CoV-2 in Ferrets. Cell Host Microbe 27, 704–709.e2.

Kochi, A.N., Tagliari, A.P., Forleo, G.B., Fassini, G.M., and Tondo, C. (2020). Cardiac and arrhythmic complications in patients with COVID-19. J. Cardiovasc. Electrophysiol. 31, 1003–1008.

Kraemer, A.M., Saraiva, L.R., and Korsching, S.I. (2008). Structural and functional diversification in the teleost S100 family of calcium-binding proteins. BMC Evol. Biol. 8, 1–23.

Ku, M.-W., Bourgine, M., Authié, P., Lopez, J., Nemirov, K., Moncoq, F., Noirat, A., Vesin, B., Nevo, F., Blanc, C., et al. (2020). Intranasal Vaccination with a Lentiviral Vector Strongly Protects against SARS-CoV-2 in Mouse and Golden Hamster Preclinical Models. BioRxiv. doi: https://doi.org/10.1101/2020.07.21.214049

Kuba, K., Imai, Y., Rao, S., Gao, H., Guo, F., Guan, B., Huan, Y., Yang, P., Zhang, Y., Deng, W., et al. (2005). A crucial role of angiotensin converting enzyme 2 (ACE2) in SARS coronavirus-induced lung injury. Nat. Med. 11, 875–879.

Kuck, K.H. (2020). Arrhythmias and sudden cardiac death in the COVID-19 pandemic. Herz 45, 325–326.

Lamers, M.M., Beumer, J., Vaart, J. Van Der, Knoops, K., Puschhof, J., Breugem, T.I., Ravelli, R.B.G., Schayck, J.P. Van Mykytyn, A.Z., Duimel, H.Q., et al. (2020). SARS-CoV-2 productively infects human gut enterocytes. Science 369, 50–54.

Levraud, J.-P., Jouneau, L., Briolat, V., Laghi, V., and Boudinot, P. (2019). IFN-Stimulated Genes in Zebrafish and Humans Define an Ancient Arsenal of Antiviral Immunity. J. Immunol. 203(12), 3361–3373

Li, L., Chong, H.C., Ng, S.Y., Kwok, K.W., Teo, Z., Tan, E.H.P., Choo, C.C., Seet, J.E., Choi, H.W., Buist, M.L., et al. (2015). Angiopoietin-like 4 increases pulmonary tissue leakiness and damage during influenza pneumonia. Cell Rep. 10, 654–663.

Libby, P. (2020). The Heart in COVID-19: Primary Target or Secondary Bystander? JACC Basic to Transl. Sci. 5, 537–542.

Liu, Z., Xiao, X., Wei, X., Li, J., Yang, J., Tan, H., Zhu, J., Zhang, Q., Wu, J., and Liu, L. (2020). Composition and divergence of coronavirus spike proteins and host ACE2 receptors predict potential intermediate hosts of SARS-CoV-2. J. Med. Virol. 92, 595–601.

Lopes, R.D., Macedo, A.V.S., Silva, P.G.M. de B. e, JunqueiraMoll-Bernardes, R., Feldman, A., Arruda, G.D.S., Souza, A.S. de, Albuquerque, D.C. de, Mazza, L., Santos, M.F., et al. (2020). Continuing versus suspending angiotensin-converting enzyme inhibitors and angiotensin receptor blockersLJ: Impact on adverse outcomes in hospitalized patients with severe acute respiratory syndrome coronavirus 2 (SARS-CoV-2) – The BRACE CORONA Trial. Am. Heart J. 226, 50–59.

Lucas, C., Wong, P., Klein, J., Castro, T.B.R., Silva, J., Sundaram, M., Ellingson, M.K., Mao, T., Oh, J.E., Israelow, B., et al. (2020). Longitudinal analyses reveal immunological misfiring in severe COVID-19. Nature 584, 463–469.

Ludwig, M., Palha, N.¤, Torhy, C., Rie Briolat, V., Colucci-Guyon, E., Bré Mont, M., Herbomel, P., Boudinot, P., and Levraud, J.-P. (2011). Whole-Body Analysis of a Viral Infection: Vascular Endothelium is a Primary Target of Infectious Hematopoietic Necrosis Virus in Zebrafish Larvae. PLoS Pathog 7(2):e1001269.

Lutz, C., Maher, L., Lee, C., and Kang, W. (2020). COVID-19 preclinical models: Human angiotensin-converting enzyme 2 transgenic mice. Hum. Genomics 14, 1–11.

Mellert, T.K., Getchell, M.L., Sparks, L., and Getchell, T. V (1992). Characterization of the immune barrier in human olfactory mucosa. Otolaryngol Head Neck Surg. 106, 181–188.

Menni, C., Valdes, A.M., Freidin, M.B., Sudre, C.H., Nguyen, L.H., Drew, D.A., Ganesh, S., Varsavsky, T., Cardoso, M.J., El-Sayed Moustafa, J.S., et al. (2020). Real-time tracking of self-reported symptoms to predict potential COVID-19. Nat. Med. 26, 1037–1040.

Mudd, P.A., Crawford, J.C., Turner, J.S., Souquette, A., Reynolds, D., Bender, D., Bosanquet, J.P., Anand, N.J., Striker, D.A., Martin, R.S., et al. (2020). Targeted Immunosuppression Distinguishes COVID-19 from Influenza in Moderate and Severe Disease. MedRxiv. doi: 10.1101/2020.05.28.20115667.

Munster, V.J., Feldmann, F., Williamson, B.N., van Doremalen, N., Pérez-Pérez, L., Schulz, J., Meade-White, K., Okumura, A., Callison, J., Brumbaugh, B., et al. (2020). Respiratory disease in rhesus macaques inoculated with SARS-CoV-2. Nature 585, 268–272.

Palha, N., Guivel-Benhassine, F., Briolat, V., Lutfalla, G., Sourisseau, M., Ellett, F., Wang, C.H., Lieschke, G.J., Herbomel, P., Schwartz, O., et al. (2013). Real-Time Whole-Body Visualization of Chikungunya Virus Infection and Host Interferon Response in Zebrafish. PLoS Pathog. 9(9), e1003619.

Papadakis, K.A., Landers, C., Prehn, J., Kouroumalis, E.A., Moreno, S.T., Gutierrez-Ramos, J.- C., Hodge, M.R., and Targan, S.R. (2003). CC Chemokine Receptor 9 Expression Defines a Subset of Peripheral Blood Lymphocytes with Mucosal T Cell Phenotype and Th1 or T-Regulatory 1 Cytokine Profile. J. Immunol. 171, 159–165.

Park, A., and Iwasaki, A. (2020). Type I and Type III Interferons – Induction, Signaling, Evasion, and Application to Combat COVID-19. Cell Host Microbe 27, 870–878.

Patel, V.B., Clarke, N., Wang, Z., Fan, D., Parajuli, N., Basu, R., Putko, B., Kassiri, Z., Turner, A.J., and Oudit, G.Y. (2014). Angiotensin II induced proteolytic cleavage of myocardial ACE2 is mediated by TACE/ADAM-17: A positive feedback mechanism in the RAS. J. Mol. Cell. Cardiol. 66, 167–176.

Peterson, J., Lin, B., Barrios-Camacho, C.M., Herrick, D.B., Holbrook, E.H., Jang, W., Coleman, J.H., and Schwob, J.E. (2019). Activating a Reserve Neural Stem Cell Population In Vitro Enables Engraftment and Multipotency after Transplantation. Stem Cell Reports 12, 680–695.

Pfaffl, M.W. (2001). A new mathematical model for relative quantification in real-time RT– PCR. Nucleic Acids Res. 29, e45.

Postlethwait, J.H., Farnsworth, D.R., and Miller, A.C. (2020). An intestinal cell type in zebrafish is the nexus for the SARS-CoV-2 receptor and the Renin Angiotensin-Aldosterone System that contributes to COVID-19 comorbidities. BioRxiv. doi: 10.1101/2020.09.01.278366.

Ribeiro-Oliveira, A., Nogueira, A.I., Pereira, R.M., Vilas Boas, W.W., Souza dos Santos, R.A., and Simões e Silva, A.C. (2008). The renin-angiotensin system and diabetes: An update. Vasc. Health Risk Manag. 4, 787–803.

Rockx, B., Kuiken, T., Herfst, S., Bestebroer, T., Lamers, M.M., Munnink, B.B.O., De Meulder, D., Van Amerongen, G., Van Den Brand, J., Okba, N.M.A., et al. (2020). Comparative pathogenesis of COVID-19, MERS, and SARS in a nonhuman primate model. Science 368, 1012–1015.

Rodriguez, S., Cao, L., Rickenbacher, G.T., Benz, E.G., Magdamo, C., Gomez, L.R., Holbrook, E.H., Albers, A.D., Gallagher, R., Westover, M.B., et al. (2020). Innate immune signaling in the olfactory epithelium reduces odorant receptor levels: modeling transient smell loss in COVID-19 patients. MedRxiv. doi: 10.1101/2020.06.14.20131128.

Sa Ribero, M., Jouvenet, N., Dreux, M., and Nisole, S. (2020). Interplay between SARS-CoV-2 and the type I interferon response. PLoS Pathog. 16, 1–22.

Saraiva, L.R., Ahuja, G., Ivandic, I., Syed, A.S., Marioni, J.C., Korsching, S.I., and Logan, D.W. Molecular and neuronal homology between the olfactory systems of zebrafish and mouse. Sci. Rep. 5, 11487.

Sepahi, A., Casadei, E., Tacchi, L., Muñoz, P., LaPatra, S.E., and Salinas, I. (2016). Tissue Microenvironments in the Nasal Epithelium of Rainbow Trout (Oncorhynchus mykiss) Define Two Distinct CD8α^+^ Cell Populations and Establish Regional Immunity. J. Immunol. 197, 4453–4463.

Sepahi, A., Kraus, A., Casadei, E., Johnston, C.A., Galindo-villegas, J., Kelly, C., García-Morenoc, D., Muñoze, P., Mulero, V., Huertas, M., et al. (2019). Olfactory sensory neurons mediate ultrarapid antiviral immune responses in a TrkA-dependent manner. Proc. Natl. Acad. Sci. U. S. A. 116, 12428–12436.

Shi, J., Wen, Z., Zhong, G., Yang, H., Wang, C., Huang, B., Liu, R., He, X., Shuai, L., Sun, Z., et al. (2020). Susceptibility of ferrets, cats, dogs, and other domesticated animals to SARS-coronavirus 2. Science 368, 1016–1020.

Slade, C.D., Reagin, K.L., Lakshmanan, H.G., Klonowski, K.D., and Watford, W.T. (2020). Placenta-specific 8 limits IFNγ production by CD4 T cells in vitro and promotes establishment of influenza-specific CD8 T cells in vivo. PLoS One 15, 1–18.

Song, E., Zhang, C., Israelow, B., Lu-Culligan, A., Prado, A.V., Skriabine, S., Lu, P., Weizman, O.-E., Liu, F., Dai, Y., et al. (2020). Neuroinvasion of SARS-CoV-2 in human and mouse brain. BioRxiv. doi: https://doi.org/10.1101/2020.06.25.169946.

Spinato, G., Fabbris, C., Polesel, J., Cazzador, D., Borsetto, D., Hopkins, C., and Paolo Boscolo-Rizzo (2020). Alterations in Smell or Taste in Mildly Symptomatic Outpatients With SARS-CoV-2 Infection. N. Engl. J. Med. 323, 2005–2011.

Su, H., Yang, M., Wan, C., Yi, L.X., Tang, F., Zhu, H.Y., Yi, F., Yang, H.C., Fogo, A.B., Nie, X., et al. (2020). Renal histopathological analysis of 26 postmortem findings of patients with COVID-19 in China. Kidney Int. 98, 219–227.

Suzuki, M., Saito, K., Min, W.P., Vladau, C., Toida, K., Itoh, H., Murakami, S. (2007). Identification of viruses in patients with postviral olfactory dysfunction. Laryngoscope 117(2), 272–277.

Tabata, S., Imai, K., Kawano, S., Ikeda, M., Kodama, T., Miyoshi, K., Obinata, H., Mimura, S., Kodera, T., Kitagaki, M., et al. (2020). Clinical characteristics of COVID-19 in 104 people with SARS-CoV-2 infection on the Diamond Princess cruise ship: a retrospective analysis. Lancet Infect. Dis. 20, 1043–1050.

Taylor, K.L., Grant, N.J., Temperley, N.D., and Patton, E.E. (2010). Small molecule screening in zebrafish: An in vivo approach to identifying new chemical tools and drug leads. Cell Commun. Signal. 8, 1–14.

Torabi, A., Mohammadbagheri, E., Akbari Dilmaghani, N., Bayat, A.H., Fathi, M., Vakili, K., Alizadeh, R., Rezaeimirghaed, O., Hajiesmaeili, M., Ramezani, M., et al. (2020). Proinflammatory Cytokines in the Olfactory Mucosa Result in COVID-19 Induced Anosmia. ACS Chem. Neurosci. 11, 1909–1913.

Van Dycke, J., Ny, A., Conceição-Neto, N., Maes, J., Hosmillo, M., Cuvry, A., Goodfellow, I., Nogueira, T.C., Verbeken, E., Matthijnssens, J., et al. (2019). A robust human norovirus replication model in zebrafish larvae. PLoS Pathog. 15, 1–21.

Ventura-Fernandes, B.H., Feitosa, N.M., Barbosa, A.P., Bomfim, C.G., Garnique, A.M.B., Gomes, F.I.F., Nakajima, R.T., Belo, M.A.A., Eto, S.F., Fernandes, D.C., et al. (2020). Zebrafish studies on the vaccine candidate to COVID-19, the Spike protein: Production of antibody and adverse reaction. BioRxiv. doi: https://doi.org/10.1101/2020.10.20.346262.

Wadman, M., Couzin-Frankel, J., Kaiser, J., and O, C.M. (2020). A rampage through the body. Science 368, 356–360.

Wagener, B.M., Hu, M., Zheng, A., Zhao, X., Che, P., Brandon, A., Anjum, N., Snapper, S., Creighton, J., Guan, J.-L., et al. (2016). Neuronal Wiskott–Aldrich syndrome protein regulates TGF-β1–mediated lung vascular permeability. FASEB J. 30, 2557–2569.

Wan, Y., Shang, J., Graham, R., Baric, R.S., and Li, F. (2020). Receptor Recognition by the Novel Coronavirus from Wuhan: an Analysis Based on Decade-Long Structural Studies of SARS Coronavirus. J. Virol. 94, 1–9.

Wang, D., Hu, B., Hu, C., Zhu, F., Liu, X., Zhang, J., Wang, B., Xiang, H., Cheng, Z., Xiong, Y., et al. (2020a). Clinical Characteristics of 138 Hospitalized Patients with 2019 Novel Coronavirus-Infected Pneumonia in Wuhan, China. JAMA - J. Am. Med. Assoc. 323, 1061–1069.

Wang, M., Cao, R., Zhang, L., Yang, X., Liu, J., Xu, M., Shi, Z., Hu, Z., Zhong, W., and Xiao, G. (2020b). Remdesivir and chloroquine effectively inhibit the recently emerged novel coronavirus (2019-nCoV) in vitro. Cell Res. 30, 269–271.

Wei, S., Zhou, J.M., Chen, X., Shah, R.N., Liu, J., Orcutt, T.M., Traver, D., Djeu, J.Y., Litman, G.W., and Yoder, J.A. (2007). The zebrafish activating immune receptor Nitr9 signals via Dap12. Immunogenetics 59, 813–821.

De Wit, E., Van Doremalen, N., Falzarano, D., and Munster, V.J. (2016). SARS and MERS: Recent insights into emerging coronaviruses. Nat. Rev. Microbiol. 14, 523–534.

Yao, Y., Yao, J., and Boström, K.I. (2019). SOX Transcription Factors in Endothelial Differentiation and Endothelial-Mesenchymal Transitions. Front. Cardiovasc. Med. 6, 1–8.

Yoo, S.K., and Huttenlocher, A. (2011). Spatiotemporal photolabeling of neutrophil trafficking during inflammation in live zebrafish. J. Leukoc. Biol. 89, 661–667.

Yu, C.R., and Wu, Y. (2017). Regeneration and rewiring of rodent olfactory sensory neurons. Exp. Neurol. 287, 395–408.

Zhang, A.J., Lee, A.C.-Y., Chu, H., Chan, J.F.-W., Fan, Z., Li, C., Liu, F., Chen, Y., Yuan, S., Poon, V.K.-M., et al. (2020). Severe Acute Respiratory Syndrome Coronavirus 2 Infects and Damages the Mature and Immature Olfactory Sensory Neurons of Hamsters. Clin. Infect. Dis. ciaa 995. https://doi.org/10.1093/cid/ciaa995.

Zheng, Y., Vertuani, S., Nyström, S., Audebert, S., Meijer, I., Tegnebratt, T., Borg, J.P., Uhlén, P., Majumdar, A., and Holmgren, L. (2009). Angiomotin-like protein 1 controls endothelial polarity and junction stability during sprouting angiogenesis. Circ. Res. 105, 260–270.

Zhou, Y., Zhou, B., Pache, L., Chang, M., Khodabakhshi, A.H., Tanaseichuk, O., Benner, C., and Chanda, S.K. (2019). Metascape provides a biologist-oriented resource for the analysis of systems-level datasets. Nat. Commun. 10, 1523.

Zhu, J., Ji, P., Pang, J., Zhong, Z., Li, H., He, C., Zhang, J., and Zhao, C. (2020). Clinical characteristics of 3062 COVID-19 patients: A meta-analysis. J. Med. Virol. 92, 1902–1914.

Ziegler, C.G.K., Allon, S.J., Nyquist, S.K., Mbano, I.M., Miao, V.N., Tzouanas, C.N., Cao, Y., Yousif, A.S., Bals, J., Hauser, B.M., et al. (2020). SARS-CoV-2 Receptor ACE2 Is an Interferon-Stimulated Gene in Human Airway Epithelial Cells and Is Detected in Specific Cell Subsets across Tissues. Cell 181, 1016–1035.e19.

Zou, J., and Secombes, C.J. (2016). The function of fish cytokines. Biology (Basel). 5.

Zou, X., Chen, K., Zou, J., Han, P., Hao, J., and Han, Z. (2020). Single-cell RNA-seq data analysis on the receptor ACE2 expression reveals the potential risk of different human organs vulnerable to 2019-nCoV infection. Front. Med. 14, 185–192.

