## Supplemental Table 1-3 for "A zebrafish model for COVID-19 recapitulates olfactory and cardiovascular pathophysiologies caused by SARS-CoV-2"

Table S1: Phylogenetic sequence analyses of vertebrate ACE2 amino acid sequences showing the percent similarity and homology. Analyses were performed in MATGAT.

|  | ***H. sapiens*** | ***M. mulatta*** | ***M. musculus*** | ***R. norvegicus*** | ***M. putorius furo*** | ***M. javanica*** | ***R. sinicus*** | ***C. dromedarius*** | ***C. familiaris*** | ***G. gallus*** | ***A. mississippiensis*** | ***X. tropicalis*** | ***D. rerio*** |
| --- | --- | --- | --- | --- | --- | --- | --- | --- | --- | --- | --- | --- | --- |
| ***H. sapiens*** |  | **82.6** | **82.1** | **82.5** | **82.6** | **84.8** | **80.7** | **83.2** | **80.5** | **65.7** | **66.1** | **59** | **58.3** |
| ***M. mulatta*** | **91.8** |  | **81.5** | **80.1** | **100** | **86.6** | **80.7** | **84.5** | **86.3** | **66.3** | **67.4** | **59.9** | **58.7** |
| ***M. musculus*** | **89.9** | **90.3** |  | **90.4** | **81.5** | **82.7** | **78.3** | **81** | **78** | **66.4** | **67.2** | **61.6** | **58.1** |
| ***R. norvegicus*** | **90.6** | **89.3** | **94.7** |  | **80.1** | **82.6** | **78** | **80.5** | **77** | **65.9** | **66.2** | **60.1** | **58.4** |
| ***M. putorius furo*** | **91.8** | **100** | **90.3** | **89.3** |  | **86.6** | **80.7** | **84.5** | **86.3** | **66.3** | **67.4** | **59.9** | **58.7** |
| ***M. javanica*** | **91.7** | **93.5** | **90.6** | **90.2** | **93.5** |  | **82.9** | **84.7** | **83.7** | **65.7** | **67.3** | **61.2** | **58.5** |
| ***R. sinicus*** | **90.3** | **89.7** | **88.9** | **88.1** | **89.7** | **90.1** |  | **81.4** | **77.5** | **65.6** | **64.3** | **60.3** | **57.3** |
| ***C. dromedarius*** | **92.8** | **92.7** | **90.4** | **89.9** | **92.7** | **92.5** | **90.1** |  | **80.2** | **66.5** | **66.5** | **58.1** | **57.3** |
| ***C. familiaris*** | **88.1** | **90.7** | **86** | **85.1** | **90.7** | **88.7** | **85.6** | **88.4** |  | **63.7** | **64.7** | **59** | **55.7** |
| ***G. gallus*** | **80.1** | **81.3** | **81.2** | **81.4** | **81.3** | **80.6** | **80.2** | **80.2** | **76.6** |  | **74.2** | **61.1** | **57.1** |
| ***A. mississippiensis*** | **81.2** | **82.1** | **81.1** | **80.9** | **82.1** | **81.1** | **80.5** | **81.2** | **76.9** | **86** |  | **60.7** | **59.3** |
| ***X. tropicalis*** | **73.8** | **74.3** | **75.3** | **74.2** | **74.3** | **74.4** | **74.5** | **73** | **70.8** | **75.4** | **74.7** |  | **57.3** |
| ***D. rerio*** | **73.2** | **73.2** | **74.1** | **74.5** | **73.2** | **73** | **73.7** | **74** | **70.4** | **74.1** | **74.7** | **72.5** |  |
| **Similarity** | | | | | | | | | | | | | |

Table S2: Sequence analyses of SARS-CoV-2 S protein-ACE2 binding amino acids in different vertebrates. Amino acids in red denote non-conserved amino acids. In green, amino acid substitutions that represent non-functional changes.

|  | **Sequence ID** | **AA Position** | | | | | | | | | | | | | | **Similarity to human (whole protein)** | **% SARS-CoV-2 Spike binding AA** |
| --- | --- | --- | --- | --- | --- | --- | --- | --- | --- | --- | --- | --- | --- | --- | --- | --- | --- |
|  |  | **27** | **28** | **30** | **31** | **34** | **38** | **41** | **42** | **82** | **329** | **353** | **354** | **355** | **357** |  |  |
| ***H. sapiens*** | Q9BYF1 | **T** | **F** | **D** | **K** | **H** | **D** | **Y** | **Q** | **M** | **E** | **K** | **G** | **D** | **R** | **100** | **100** |
| ***M. mulatta*** | NP_001129168.1 | **T** | **F** | **E** | **K** | **Y** | **E** | **Y** | **Q** | **T** | **Q** | **K** | **R** | **D** | **R** | **82** | **71/78** |
| ***M. musculus*** | Q8R0I0 | **T** | **F** | **N** | **N** | **Q** | **D** | **Y** | **Q** | **S** | **A** | **H** | **G** | **D** | **R** | **82** | **57/71** |
| ***R. norvegicus*** | Q5EGZ1 | **S** | **F** | **N** | **K** | **Q** | **D** | **Y** | **Q** | **N** | **T** | **H** | **G** | **D** | **R** | **82** | **57/71** |
| ***M. putorius furo*** | BAE53380.1 | **T** | **F** | **E** | **K** | **Y** | **E** | **Y** | **Q** | **T** | **Q** | **K** | **R** | **D** | **R** | **82** | **57/71** |
| ***M. javanica*** | XP_017505752.1 | **T** | **F** | **E** | **K** | **S** | **E** | **Y** | **Q** | **N** | **E** | **K** | **H** | **D** | **R** | **84** | **64/78** |
| ***R. sinicus*** | U5WHY8 | **M** | **F** | **D** | **K** | **T** | **D** | **H** | **Q** | **N** | **N** | **K** | **G** | **D** | **R** | **80** | **64** |
| ***C. dromedarius*** | XP_010991717.1 | **T** | **F** | **E** | **E** | **H** | **D** | **Y** | **Q** | **T** | **D** | **K** | **G** | **D** | **R** | **83** | **64/78** |
| ***C. familiaris*** | F1P7C5 | **T** | **F** | **E** | **K** | **Y** | **E** | **Y** | **Q** | **T** | **E** | **K** | **G** | **D** | **R** | **84** | **71/86** |
| ***G. gallus*** | F1NHR4 | **T** | **F** | **A** | **E** | **V** | **D** | **Y** | **E** | **R** | **T** | **K** | **N** | **D** | **R** | **65** | **57** |
| ***A. mississippiensis*** | KYO30243.1 | **T** | **F** | **N** | **Q** | **Q** | **G** | **Y** | **E** | **K** | **I** | **M** | **K** | **D** | **R** | **66** | **35** |
| ***X. tropicalis*** | F6PSC4 | **D** | **F** | **K** | **R** | **Q** | **V** | **H** | **Q** | **A** | **N** | **M** | **N** | **D** | **R** | **58** | **35/43** |
| ***D. rerio*** | XP_005169417.1 | **E** | **F** | **N** | **K** | **E** | **D** | **Y** | **Q** | **A** | **N** | **R** | **K** | **D** | **R** | **57** | **50/64** |

Table S3: List of primers used in the present study.

| **Target** | **forward 5' --> 3'** | **reverse 5' --> 3'** | **use and comments** |
| --- | --- | --- | --- |
| ***ace2*** | CTGGCTCCTGCTTTTGGC | TCTTTATCTGCATTTTCCTGGGAG | qPCR, Tm = 62 |
| ***mxa*** | TTGACCTCCCTGGCATTGCA | GATTGTCTCTTGCCTTGTAACA | qPCR, Tm = 60 |
| ***il4/13b*** | GCAGGAATGGCTTTGAAGGGTAAA | AAACTCCTTCATTGTGCATTCCCC | qPCR, Tm = 60 |
| ***il22*** | TTGGAATCAGACGAGCACAC | GGCCAAATCCATAATTGCAC | qPCR, Tm = 60 |
| ***il17a/f3*** | AAGATGTTCTGGTGTGAAGAAGTG | ACCCAAGCTGTCTTTCTTTGAC | qPCR, Tm = 60 |
| ***ifnΦ1*** | CGCAAAGCCAGCACACAAGGA | CTCCGGATCTGCTCCCATGCT | qPCR, Tm = 60 |
| ***ifnΦ3*** | CGAGGATCAGGTTACTGGTGT | GTTCATGATGCATGTGCTGTA | qPCR, Tm = 60 |
| ***il1b*** | CGCTCCACATCTCGTACTCA | ATACGCGGTGCTGATAAACC | qPCR, Tm = 62 |
| ***ccl20a.3*** | TGATGGTGCTGACAATCGTG | CTTTGGACGGGTCTGTGCA | qPCR, Tm = 62 |
| ***tnfa*** | GCGCTTTTCTGAATCCTACG | TGCCCAGTCTGTCTCCTTCT | qPCR, Tm = 62 |
| ***rps11*** | CCCAGAGAAGCTATTGAT | TCACATCCCTGAAGCATG | qPCR, Tm = 62 |
| ***ccl19a.1*** | GCCCACGTGATGCTGTAATA | ACAGCGTCTCTCGATGAACC | qPCR, Tm = 62 |
