## Supplementary figures and images for "A zebrafish model for COVID-19 recapitulates olfactory and cardiovascular pathophysiologies caused by SARS-CoV-2"

### Figure S1

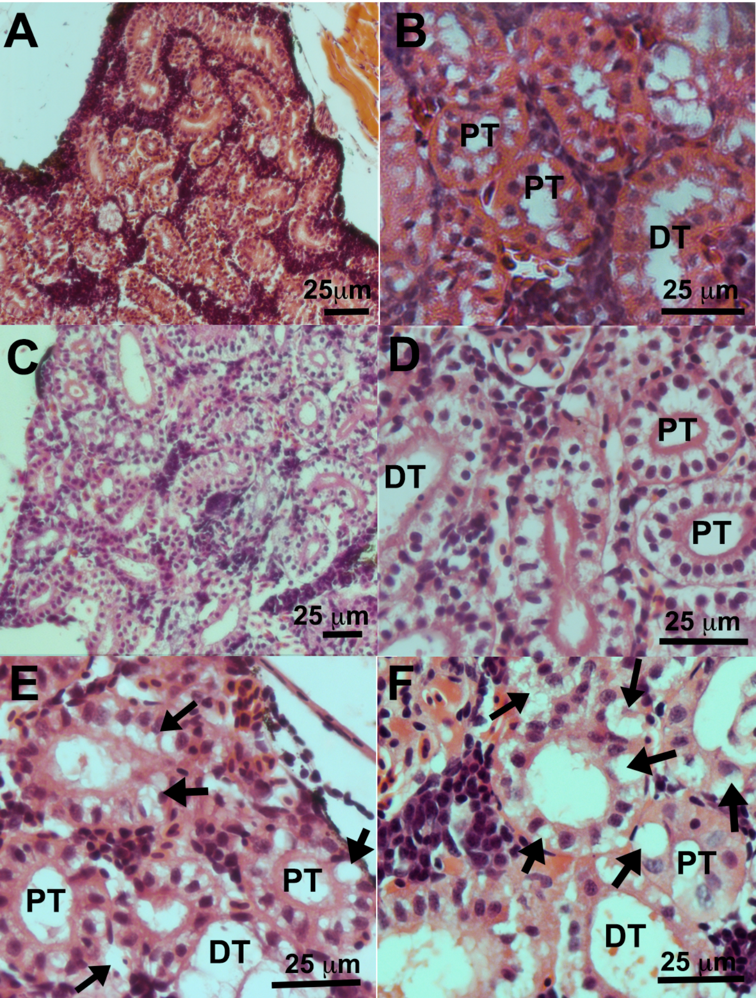

### Figure S3

#### Figure S3

A

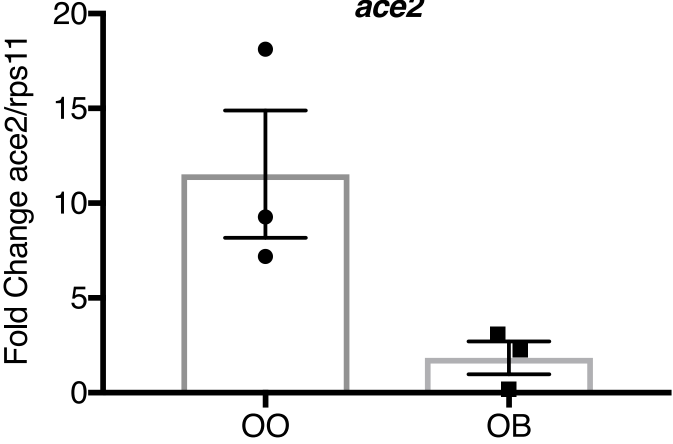
