## Supplementary material for "A zebrafish model for COVID-19 recapitulates olfactory and cardiovascular pathophysiologies caused by SARS-CoV-2": Figure S2

**A**

| | Mean Reads per Cell | Median Genes per Cell | Reads Mapped Confidently to Genome | Total Genes Detected | Median UMI Counts per Cell | Post Filtering # of Cells |
|---|---|---|---|---|---|---|
| 3 Day | 20,512 | 927 | 84.10% | 24,301 | 2,266 | 6097 |
| 3 Hour | 33,309 | 1,055 | 83.80% | 23,660 | 2,668 | 4842 |
| Control | 47,161 | 814 | 82.60% | 23,155 | 1,747 | 3613 |

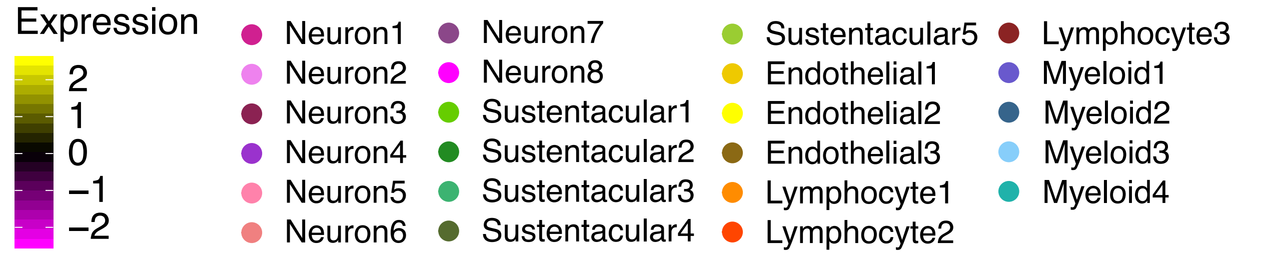


**B**

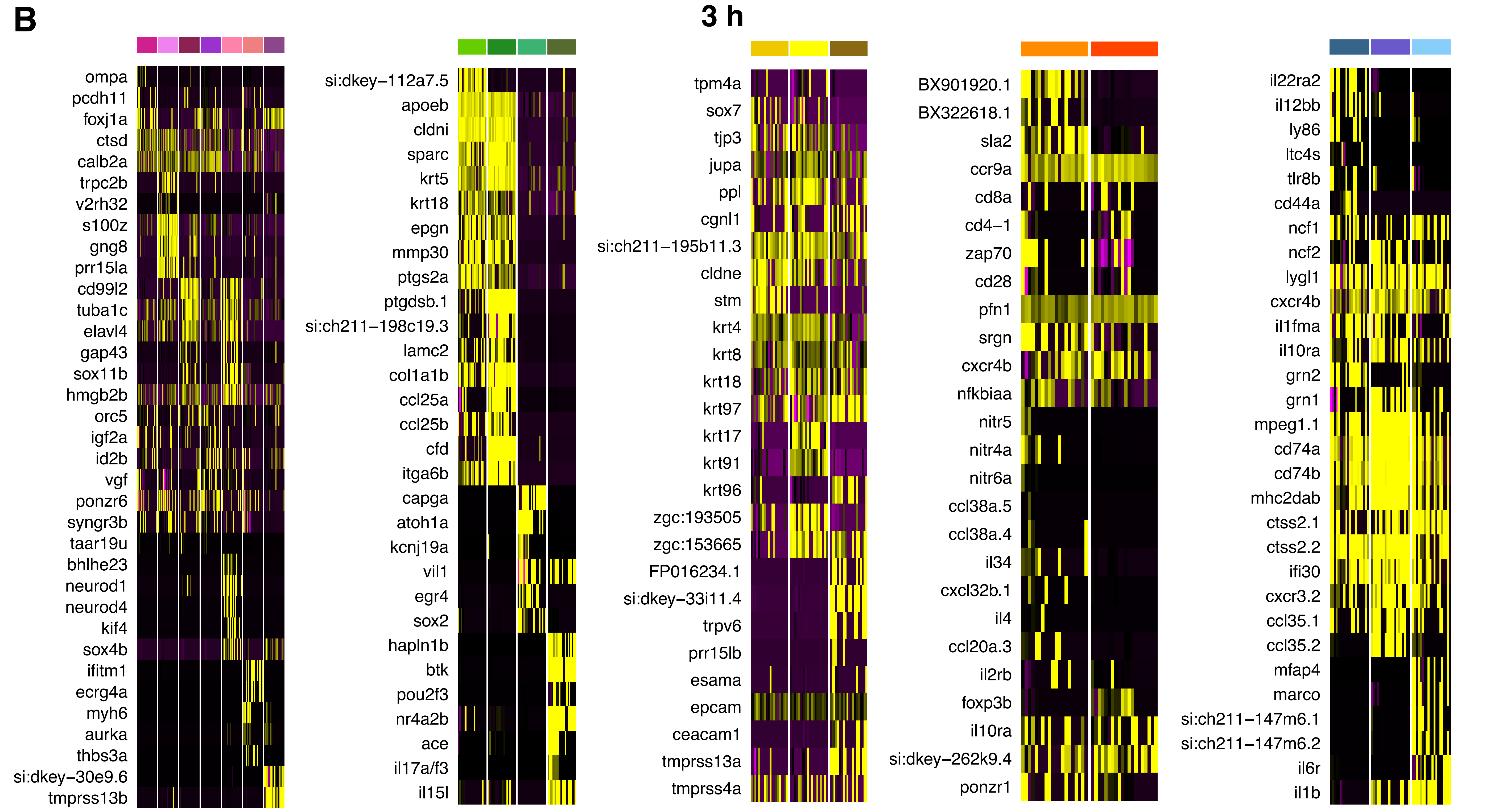


**C**

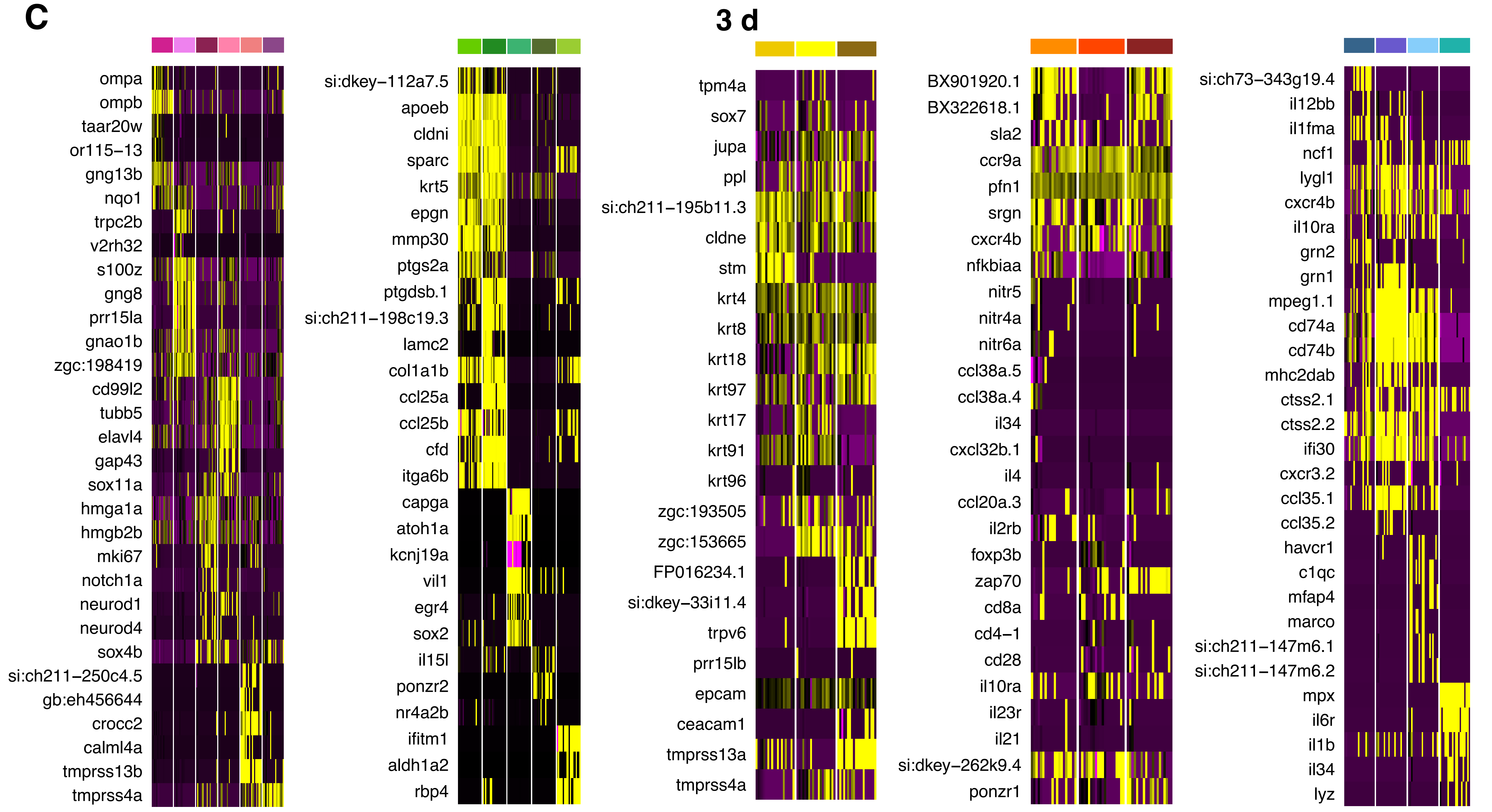
